# Loss of flavonol 3-*O*-glucosyltransferase activity confers soybean resistance to leaf-chewing insects

**DOI:** 10.1101/2025.10.01.679769

**Authors:** P.K. Prabhakar, M.A. Ortega, B-K. Ha, P.R. LaFayette, S.A. Harding, C.J. Tsai, B.R. Urbanowicz, H.R. Boerma, W.A. Parrott

## Abstract

Caterpillars and beetles are among the most economically damaging defoliating insects, and their economic damage is predicted to increase in the coming decades. Hence the use of genetically derived resistance to supplement other pest control strategies is warranted. In soybean (*Glycine max* (L.) Merr.), a major determinant for resistance is the quantitative trait locus, QTL-M. *Glyma07g14530*, the gene underlying QTL-M, encodes a feeding-inducible flavonol 3-*O*-glycosyltransferase (F3GlcT or UGT78D2) that glucosylates kaempferol, as well as quercetin, myricetin, and isorhamnetin. The resistant allele has a premature stop codon in it, thus preventing the glucosylation and sequestration of flavonols in the vacuole, leading to a concomitant accumulation of proanthocyanidins and manifestation of resistance. Expressing the dominant (susceptible) allele in resistant plants restores susceptibility and silencing the susceptible allele results in resistance. The discovery and characterization of *GmF3GlcT* helps clarify the role of flavonoids in resistance to leaf-chewing insects and facilitates the development of insect-resistant cultivars that ultimately can lower production costs and reduce insecticide applications.

**Graphical Abstract:** 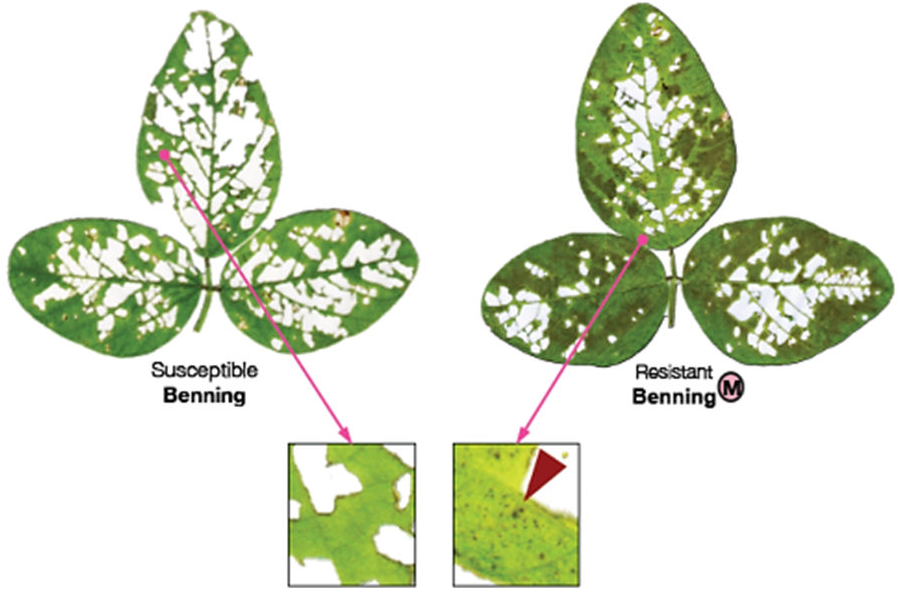

The loss of flavonol 3-O-glucosyltransferase in soybean reduces feeding damage from defoliating insects and is accompanied by a concomitant increase in proanthocyanidins (red arrowhead).

**Significance:** Unraveling the biochemical and genetic basis of soybean resistance to leaf-chewing insects facilitates the development of new, naturally insect resistant varieties. Such varieties contribute to on-farm profitability and reduced concerns over pesticide residues in the field.

## Introduction

Effective, economical, and environmentally safe pest control measures are necessary to protect crops of the future (Deutsch et al., 2018, Skendzic et al., 2021). One strategy that is immediately available is the direct deployment through breeding of host plant insect resistance (Smith and Clement, 2012) into crop plants. Case in point, soybean [*Glycine max* (L.) *Merr*] is among the world’s main sources of vegetable oil and is the main source of vegetable protein (Pagano and Miransari, 2016). Yet, 9% of worldwide soybean production is lost to pests, including insects (Oerke, 2006). In the US alone, soybean losses to leaf defoliators and their control costed almost $390 million in 2023 (Crop-Protection-Network). At the same time, current chemical-based control strategies are losing effectiveness, while the amount of insecticide applied to soybeans in the USA quadrupled between 2002 and 2012.

As an alternative to insecticides, there are soybean genotypes with a measure of resistance to defoliating insects (**Fig 1**). The Japanese landrace, ‘Sodendaizu’ (PI 229358), was one of the first reported to be insect-resistant (Van Duyn et al., 1971). The main basis for its resistance is due to a major QTL, known as QTL-M (Rector et al., 2000a, Rector et al., 2000b). Narvel et al. (2001) found QTL-M in 13 insect-resistant lines developed by different breeding programs, establishing that QTL-M is effective in different genetic backgrounds, and it continues to be broadly deployed for insect resistance in breeding programs.

**Fig 1.**
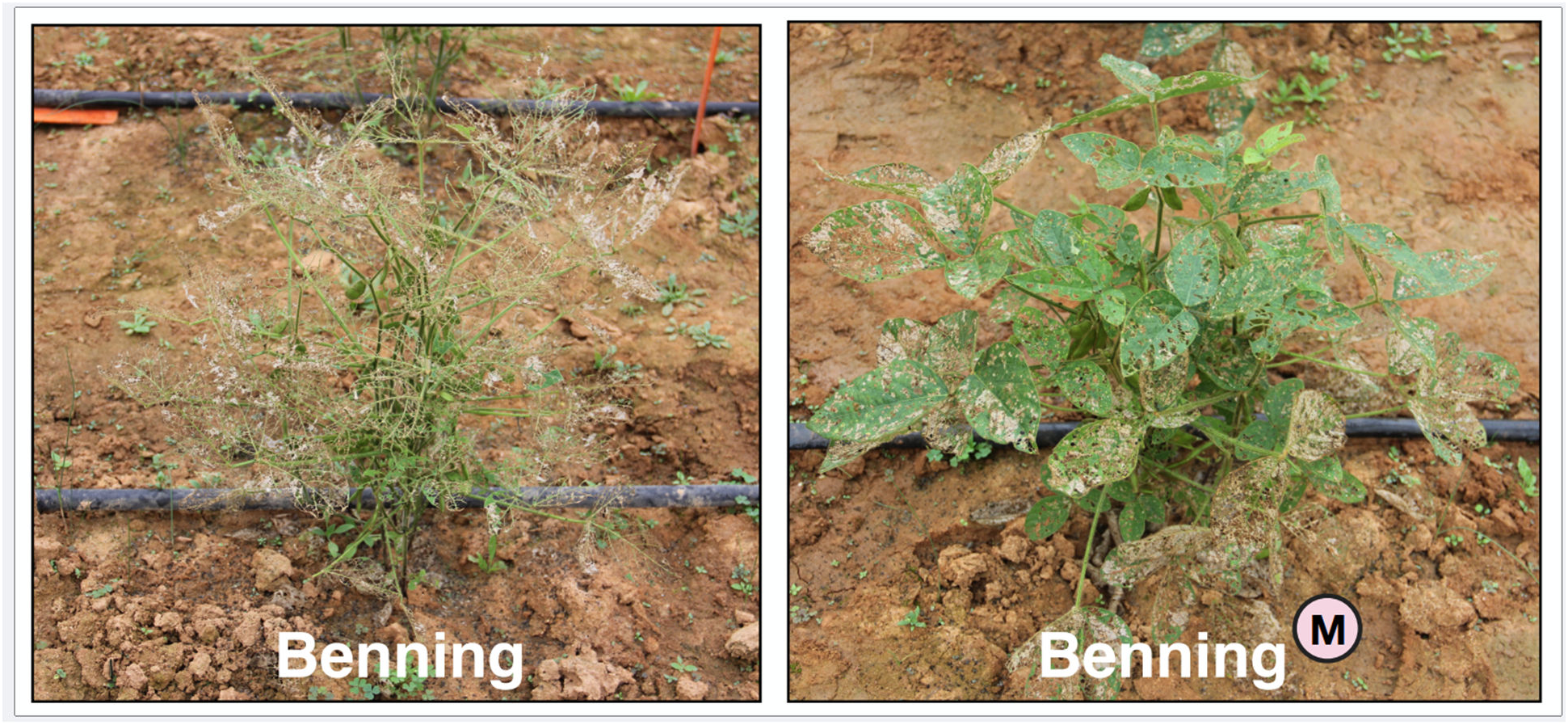
QTL-M confers resistance to leaf-chewing insects in soybean. The susceptible cultivar Benning and its insect-resistant near isoline Benning^M^ exposed to soybean looper (SBL) caterpillars in a field-cage assay. Benning^M^ was derived by backcrossing the QTL-M region into it.

QTL-M is unusual in that it provides broad resistance to multiple insects, including coleopterans, lepidopterans, pentatomid hemipterans, and even thysanopterans (Ribeiro et al., 2025). Importantly, the presence of QTL-M does not compromise feed quality or safety of soybean meal (Ortega et al., 2016b). QTL-M also enhances insect resistance when pyramided with Bt transgenes and when combined with QTL-E, another QTL conferring insect resistance (Ortega et al., 2016a). Currently, ‘G17-5173R2,’ a breeding line that combines QTLs-M and -E, has been the top yielder in replicated regional field trials (Gillen, 2021, Mailhot, 2022, Gillen, 2023).

We previously identified the causal gene underlying QTL-M as *Glyma07g14530* (Ortega, 2016, Ortega et al., 2017), which has since been validated by another research team (Zhang et al., 2022). Here we describe the identification of *Glyma07g14530* as the causal gene underlying QTL-M.

The protein encoded by *Glyma07g14530* is a family 1 glycosyltransferase (GT1), often referred to as UDP-dependent glycosyltransferases (UGTs), and found that the resistant allele contains a premature stop codon. This family of enzymes are known to play crucial roles in plant metabolism by catalyzing the transfer of a sugar molecule from and UDP-sugar donor onto a variety of small acceptor molecules, modifying their chemical and functional properties. Biochemical analysis of protein encoded by *Glyma07g14530,* showed that it is a selective flavonol 3-*O*-glucosyltransferase. Taken together, our data present a defense mechanism where plants with a functional copy of this GT render soybean plants susceptible to leaf chewing insects by glucosylating and sequestering defensive flavonols like kaempferol and quercetin. In contrast, non-glycosylated defense molecules accumulate in resistant plants leading to the observed pest-deterring trait.

Knowledge of the biochemical basis for resistance conferred by QTL-M will help design and rapidly deploy economically viable insect resistance management strategies in the longer term.

## Results

### Fine mapping and sequencing of the QTL-M region

Genetic mapping placed QTL-M in a 0.52-cM interval (1.7 Mb) on its namesake Linkage Group M, now chromosome 7 (Zhu et al., 2006). Recombinant substitution lines (RSLs), containing different PI 229358 introgressions of QTL-M into the susceptible ‘Benning’ background (**Fig S1A**), were developed to fine-map QTL-M to an interval suitable for cloning. Insect resistance (**Fig S1B**) co-segregated with a 182-kb segment (**Fig S1C**). In Williams 82, this segment contains 11 annotated gene models, none of which resembled any canonical gene involved in plant immunity known at the time. The differences between PI 229358 and Williams 82 consisted of 216 SNPs and 68 indels, including a 5.4-kb insertion in PI 229358, and 588-bp and 599-bp deletions in PI 229358. Seven gene models had polymorphic exon sequences and therefore considered candidate genes (**Table S1**).

### Identification ofh candidate insect resistance genes in QTL-M

To narrow the number of candidate genes, the seven gene models polymorphic between PI 229358 and Williams 82 were cloned from 34 insect-susceptible soybean accessions and from ‘Benning’ and ‘Jack’ (**Fig S2A**). Gene models from PI 229358 that had an allele in common with any susceptible accession were excluded as candidate genes. Only two gene models, *Glyma0714470*, a predicted P-loop-NTPase, and *Glyma07g14530,* a predicted isoflavone glycosyltransferase, have SNP alleles unique to PI 229358 (**Fig S2B**). RT-PCR indicated both genes were expressed in soybean leaves before insect attack, but only *Glyma07g14530* is upregulated after insect attack in both the susceptible Benning and the resistant Benning^M^ (**Fig 2A**) a near isogenic line of Benning containing the resistant allele of QTL-M (Fig 2A). The full-length cDNA of *Glyma07g14470* showed that contrary to the original annotation, its SNP is inside an intron (**Fig 2B**). *Glyma07g14530* consists of a single 1476-bp exon; thus, the original gene model (Glyma 1.0) was also mis-annotated. In the corrected annotation, the unique SNP is in an exon.

**Fig. 2.**
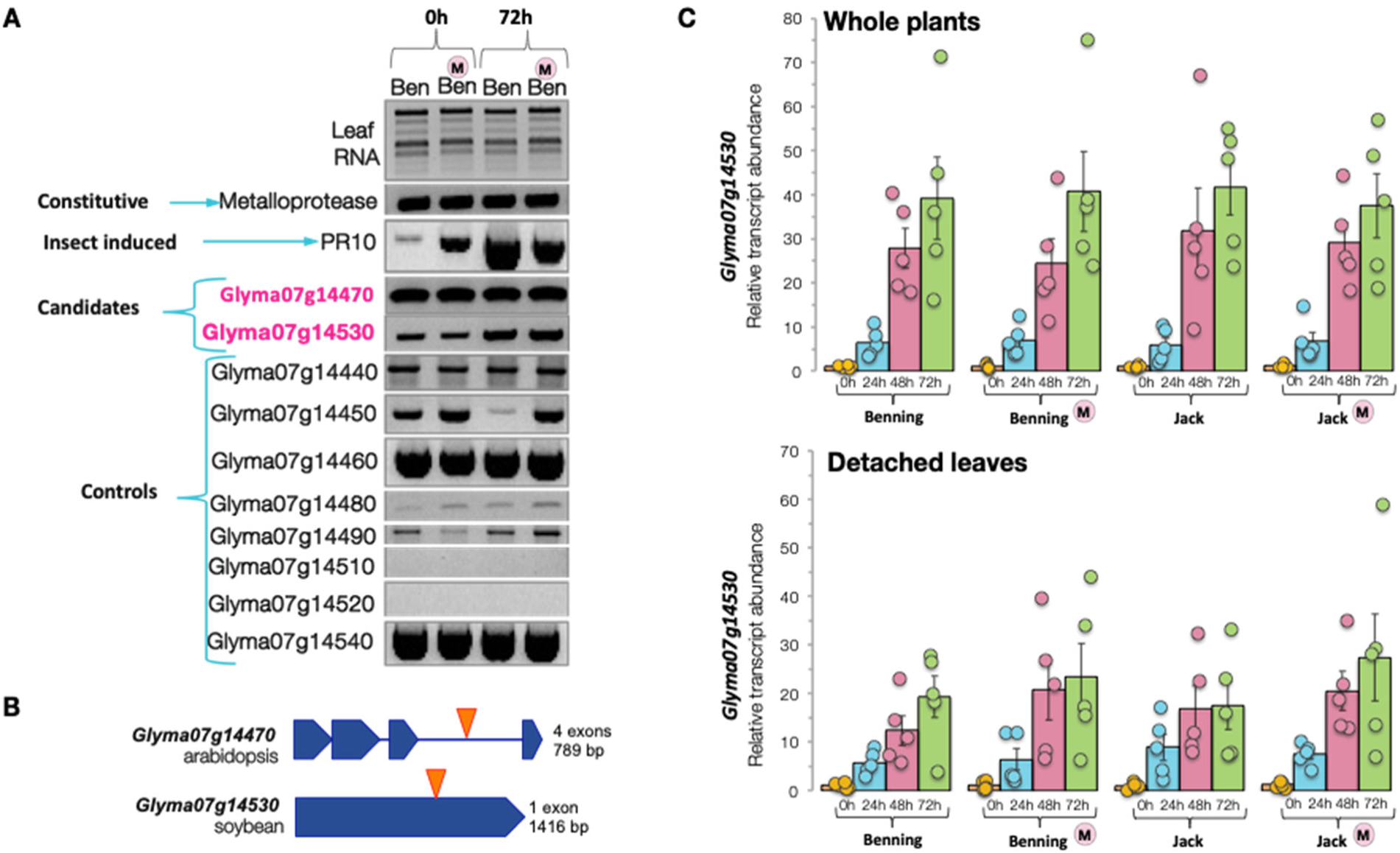
Molecular characterization of QTL-M candidate genes. (**A**) Expression of the candidate genes from the QTL-M region in leaf tissue. RNA isolated from Benning and Benning^M^ plants, at 0 hr and 72 hr after infestation with SBL caterpillars, was used as template for RT-PCR reactions. Metalloprotease, PR10, and neighboring genes in the QTL-M region were used as controls. (**B**) Full-length *Glyma07g14470* and *Glyma07g14530* transcripts. Sequences were assembled from 5’-RACE and 3’-RACE products. (**C**) Relative expression of *Glyma07g14530*. Results from infested whole plants and detached leaves are shown in the top and bottom panels respectively. RNA isolated from Benning, Benning^M^, Jack, and Jack^M^ (Jack with QTL-M backcrossed into it) at 0, 24, 48, and 72 hr after infestation was used as template for qRT-PCR. Bars represent means ± SEM from five biological replicates.

To further confirm that *Glyma07g14530* is the QTL-M causal gene, both *Glyma07g14470* and *Glyma07g14530* were cloned from a panel of 17 cultivated and wild soybean (*Glycine soja* Sieb. & Zucc.) accessions reported to be insect resistant (**Fig S2C**). The PI 229358 allele for *Glyma07g14470*’s SNP was shared with six resistant accessions, including PI 227687, which has resistance that is not associated with QTL-M (Rector et al., 2000a, Rector et al., 2000b). In contrast, the PI 229358 allele for *Glyma07g14530* is shared with only four other resistant accessions, but not with PI 227687 (**Fig S2D**).

### The resistant *Glyma07g14530* allele has a premature stop codon

The wild-type *Glyma0714530* protein (termed P530S) in susceptible soybeans is 491 amino acids long. The resistant SNP allele (cDNA position 825) results in a nonsense mutation of TG**G** (^275^W) to TG**A** (stop) that leads to a truncated protein (termed Glyma530R) of 274 amino acids that lacks the entire C-terminal Rossmann-like domain that harbors catalytic residues involved in UDP-glucose binding (magenta color in **Fig 3**).

**Fig 3.**
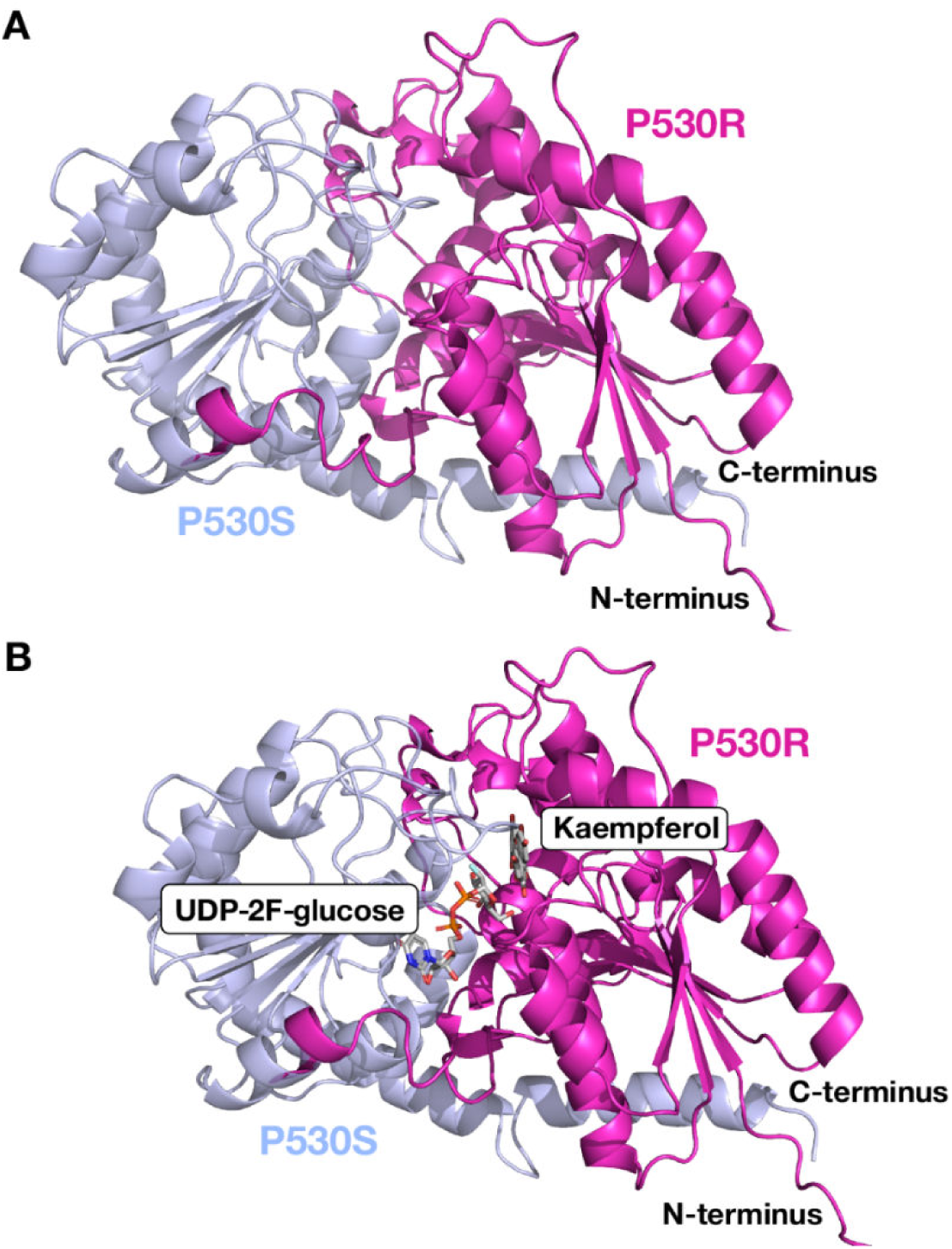
Structural comparison of full-length Glyma530S (slate) and its truncated version Glyma530R (magenta) from soybean. AlphaFold 3 was utilized to generate a predictive protein model that is shown rendered using PyMOL (DeLano Scientific LLC). (a) The single-nucleotide polymorphism (SNP) in P530-R is a nonsense mutation encoding a stop codon in place of a sense codon at residue 275 resulting in synthesis of a truncated protein (P530S, magenta) lacking key catalytic and substrate binding residues relative to the full-length counterpart (530R, lavender). (b) P530S and P530R aligned with anthocyanidin 3-*O*-glucosyltransferase in complex with Kaempferol from *Clitoria ternatea* L. (4REL; PDB id, RMSD is 1.63Ǻ using 292 Cα atoms, protein is hidden), and *Phytolacca americana* L. uridine diphosphate glycosyltransferase with Kaempferol and UDP-2-fluoro-glucose (7VEJ; PDB id, RMSD is 1.57Ǻ using 264 Cα atoms, protein is hidden). The P530R truncated protein lacks the nucleotide sugar donor binding site that is critical for activity.

The *Glyma07g14530* promoter contains four W-box motifs (TTGAC), which are predicted binding sites for WRKY transcription factors that regulate plant immune responses to stress (Eulgem and Somssich, 2007). Therefore, the hypothesis was that resistance is achieved by the loss-of-function of *Glyma07g14530,* whereas susceptibility is conferred by the functional gene. Consistent with this hypothesis, QTL-M resistance is inherited as a partially recessive locus, while plants heterozygous for QTL-M have an intermediate level of resistance (Rector et al., 2000a).

Overexpression and gene-silencing were used to test the loss-of-function hypothesis. T_2_ complementation lines expressing either transgenic or native *Glyma07g14530-S* (i.e., the wild-type allele for susceptibility) were evaluated for resistance in choice bioassays with soybean looper (SBL, *Chrysodeixis includens* Walker) caterpillars (**Fig 4**). The same was done for lines with *Glyma07g14530-S* silenced. Jack^M^ (cultivar Jack with the *Glyma07g14530-R* allele backcrossed into it) plants expressing transgenic *Glyma07g14530-S* were more defoliated than plants of Jack^M^, but less defoliated than plants of Jack (**Fig 4A**).

**Fig. 4.**
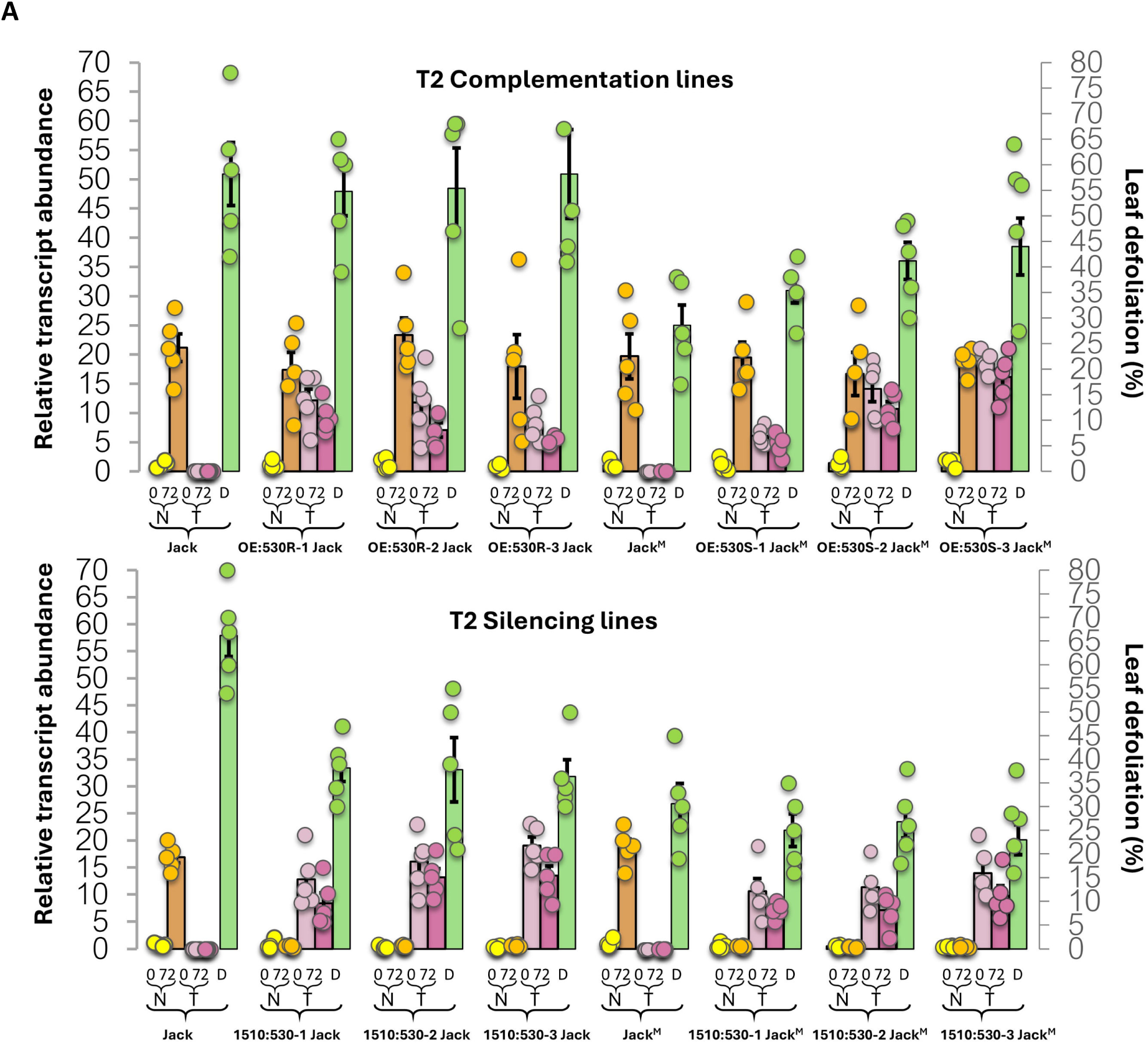
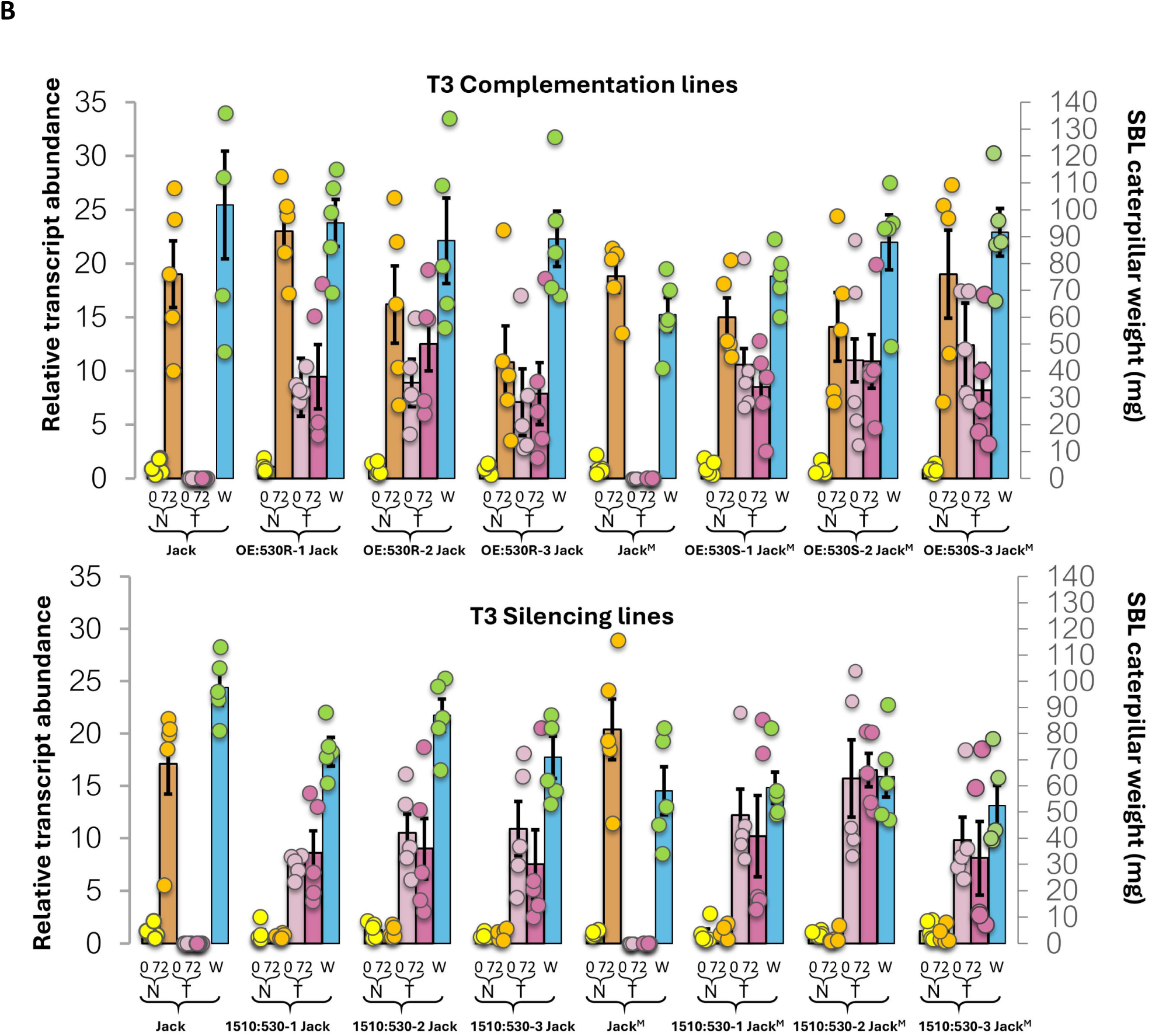
Characterization of T_2_ overexpression and silencing lines for defoliation (A) and caterpillar weight (B). The first two columns are relative transcript abundance of the *Glyma07g14530* resistant (530R) or susceptible (530S) native (N) allele at 0 and 72 h post-infestation. The second two columns are the same measure for the transgenic (T) allele or silencing construct. The fifth column represents % defoliation (D) or weight in mg (W). Bars represent means ± SEM from five biological replicates.

In the T_3_ generation no-choice assays, caterpillars feeding on Jack^M^ plants expressing *Glyma07g14530-S* were larger than caterpillars feeding on Jack^M^; two of the lines produced caterpillars that were as large as those fed on Jack. Those feeding on the silencing lines (which exhibited increased resistance) were significantly smaller. In contrast, lines of Jack with the susceptibility gene silenced were less defoliated than Jack controls; i.e., silencing of the susceptibility gene increased resistance (**Fig 4B & Fig. S4**).

*Glyma07g14530* is induced in whole plants as early as 24-hr after infestation with caterpillar neonates; at 72-hr post infestation, *Glyma07g14530* transcripts are up to 40-fold higher than those in non-infested leaves. Even in detached leaves, a 27-fold higher induction took place. No significant differences in induction level were found between resistant and susceptible RILs, or between Jack and Benning, which are both susceptible, indicating that resistance is not caused by differences in transcription levels (**Fig 2C**). By implication, defoliator feeding increases susceptibility to further defoliation by upregulating the susceptible allele of *Glyma07g14530*.

### Glyma07g14530 is a flavonol 3-O-glucosyltransferase

The full-length, wild-type *Glyma07g14530-S* was successfully expressed in *Escherichia coli* as an maltose-binding protein (MBP) fusion protein. The full-length protein is referred to here as P530S, whereas P530R refers to the truncated protein from the *Glyma07g14530* allele that lacks a complete catalytic site, and which confers resistance. Analysis of the purified P530S protein by SDS-PAGE showed that it matches the theoretical molecular weight of 97.5 kDa (**Fig 5**). Since there was no initial information on the biochemical properties of this enzyme, we developed a high-throughput UDP-Glo^TM^-based glycosyltransferase assay to determine acceptor and NDP-sugar donor specificity. Screening of acceptor substrates was done in pooled groups of four using UDP-glucose as a donor (**Fig S6**). The highest activity was found in two acceptor groups containing the following flavonoids: apigenin, kaempferol, myricetin, isorhamnetin, daidzein, genistein, gallocatechin, epigallocatechin. Further analysis of individual acceptors determined the enzyme preferred kaempferol (**Fig 5A**).

**Fig. 5.**
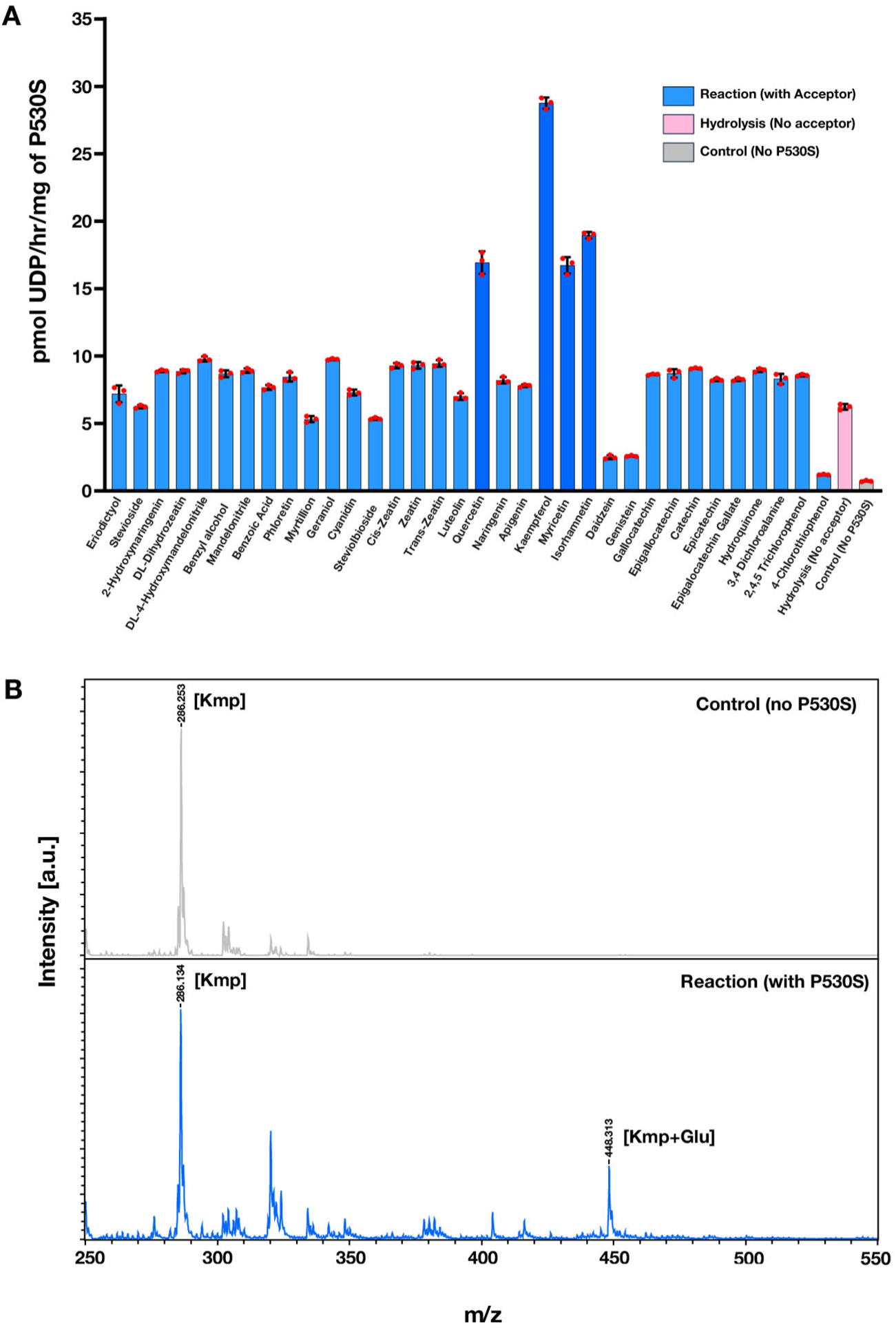
Biochemical analysis of P530S to identify acceptor substrate selectivity. (A) P530S glycosyltransferase enzyme activities were performed with 100 µM of UDP-Glc donor and 50 µM of each acceptor for 120 min. Transfer to water (hydrolysis) was measured in the absence of an acceptor. The flavonols preferred by P530 are highlighted in dark blue. Activity is represented as the mean ± standard deviation for three technical replicates (red circles). (b) MALDI-TOF MS of the products after a 16-hour reaction in the absence and presence of P530S. The series annotated [M^+^H]^+^ ions are due to structures with a mass difference of 162 Da, consistent with the sequential addition of a Glc residue to kaempferol (Kmp).

Thus, kaempferol was used to determine the pH (**Fig S7a**) and temperature (**Fig S7b**) optima and the NDP-donor selectivity. Of the seven UDP-sugars tested, UDP-Glc showed the highest hydrolytic activity (**Fig 6**). Using the optimized conditions established, 34 various acceptor substrates were screened for glucosyltransferase activity. The flavonols quercetin, myricetin, and isorhamnetin showed 65% of the activity of kaempferol when used as acceptor substrates, while no activity was present when using any of the flavanol, flavanone, or anthocyanidin acceptors. P530S was unable to use 2,4,5-trichlorophenol (TCP), 3,4-dichloroaniline (DCA), and 4-chlorothiophenol (CTP), which are common acceptors used to test O- and N-glycosylation. Matrix-Assisted Laser Desorption/Ionization-Time of Flight mass spectrometry (MALDI-TOF MS) analysis of reaction products generated when using UDP-Glc as a donor and kaempferol as an acceptor shows a product produced with a mass difference of 162 Da, consistent with the addition of a glucosyl residue to the acceptor, confirming P530S catalyzes the transfer of glucose to kaempferol (**Fig 5B**). More specifically, it is a *flavonol 3-O-glucosyltransferase*, *GmF3GlcT* for short, and enzyme-catalyzed glucosylation of target flavanols results in susceptibility to insect feeding.

**Fig 6.**
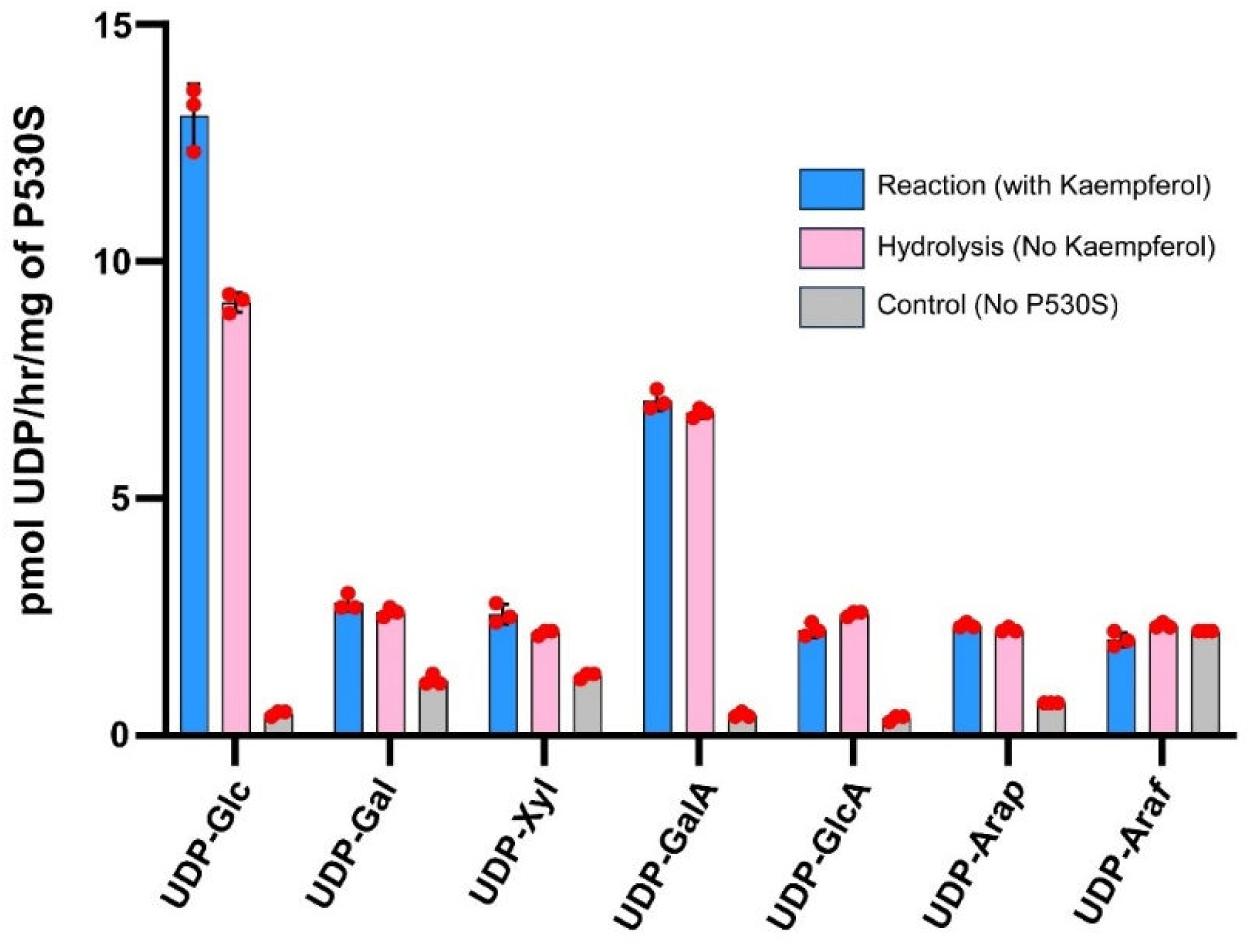
Identification of the UDP-sugar donor for P530S. Glycosyltransferase enzyme activities were performed with 50 µM of Kaempferol for 120 min with 100 µM of the indicated UDP-sugar. Transfer to water (hydrolysis) was measured in the absence of acceptor. Control represents the background without the enzyme, as UDP can be present in some NDP-sugar preparations. Activity is represented as the mean ± standard deviation for n =3 technical replicates (red circles).

### Flavonoid metabolites

A flavonol-glucoside conjugate with exact mass corresponding to that of kaempferol 3-O-β-D-glucosyl-(1-2)-β-D-glucosyl-(1-2)-β-D-glucoside is lower in both Benning^M^ and Jack^M^ before and during insect feeding. This conjugate, which could result from sequential glucosylations (Trapero et al., 2012, Yonekura-Sakakibara et al., 2019), is clearly reduced in the resistant lines (**Fig 7B**). This observation is consistent with the previous finding of flavonol 3-*O*-triglycosides in insect-resistant (Murai et al., 2013) and susceptible genotypes of soybean (Lee et al., 2022). Dillon et al. (2017) reported the synthesis and their negative impact on caterpillars of two quercetin triglycosides and a kaempferol triglycoside (presumably not triglucosides) glucose and galactose produced by soybean in the presence of UV light.

**Fig 7.**
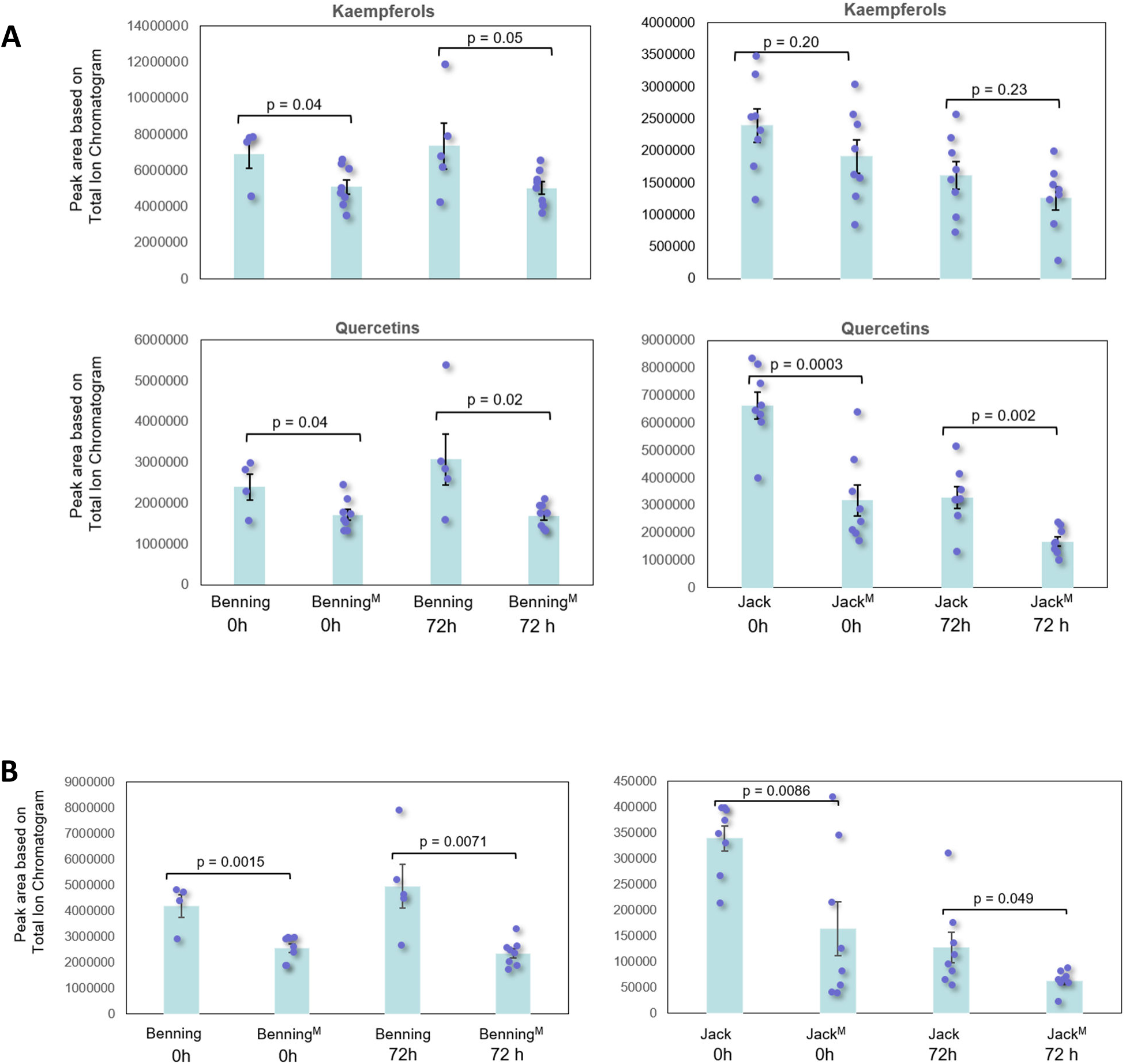
The most abundant flavonol glycosides in Benning, Benning^M^, Jack, and Jack^M^, at 0 and 72 hr after SBL infestation. (A) Kaempferols measured are Kaempferol 3-O-β-D-glucosyl-(1-2)-β-D-glucosyl-(1-2)-β-D-glucoside, Kaempferol 3-gentiobioside 7-rhamnoside, and Kaempferol 3-rhamnoside 7-galacturonide. Quercetins measured are Quercetin 3,7-dirhamnoside, Quercetin 3-O-(6“-malonyl-glucoside) 7-O-glucoside, Quercetin 3-sophoroside, Quercetin 7-glucuronide 3-rhamnoside, and Rutin. Kaempferol glycosides predominate in Benning while quercetin glycosides predominate in Jack. (B) Kaempferol 3-O-β-D-glucosyl-(1-2)-β-D-glucosyl-(1-2)-β-D-glucoside, one of the metabolites present, is lower in the resistant lines, both before and after feeding by SBL. Bars represent means ± SEM from five biological replicates.

### Proanthocyanidin precursors accumulate upon insect feeding

The levels of proanthocyanidins (PA) were compared between susceptible and QTL-M plants, because red/brown spots that are consistent with PA accumulation develop on the leaves of QTL-M plants after infestation (**Fig 8a**). PA levels increased in both Benning^M^ and Jack^M^ after feeding, while they remain constant in Benning and Jack (**Fig 8b**).

**Fig 8.**
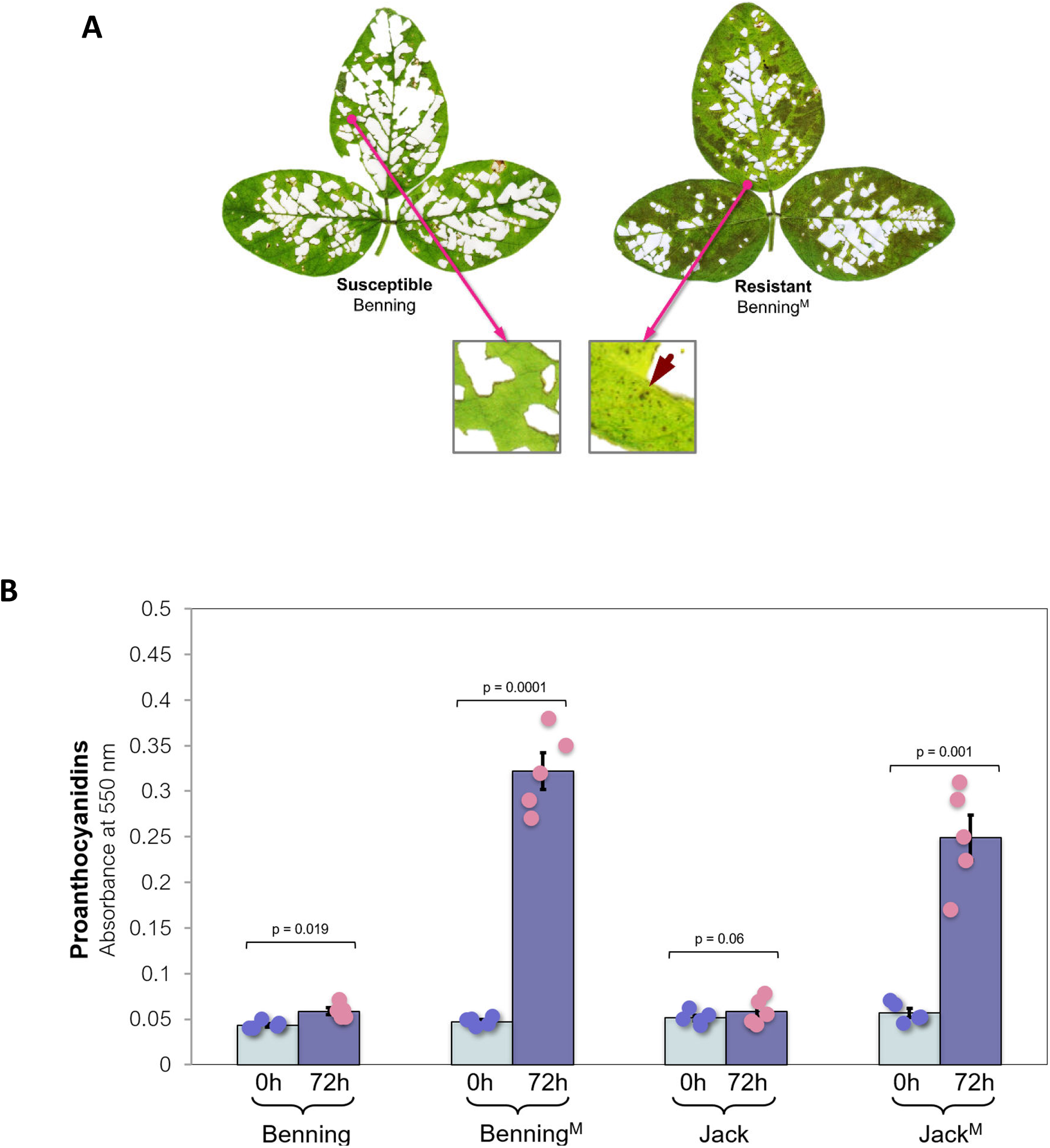
Proanthocyanidins (PA) in Benning, Benning^M^, Jack, and Jack^M^, at 0 and 72 hr after SBL infestation. **(A)** Leaves from Benning and Benning^M^ collected 10 days after SBL infestation. **(B)** PA precursors leaf tissue. The superscript “M” indicates that the genotype contains the resistant allele of QTL-M backcrossed into it. Note the increase in PAs in the resistant lines. Bars represent means ± SEM from five biological replicates.

**Fig 9.**
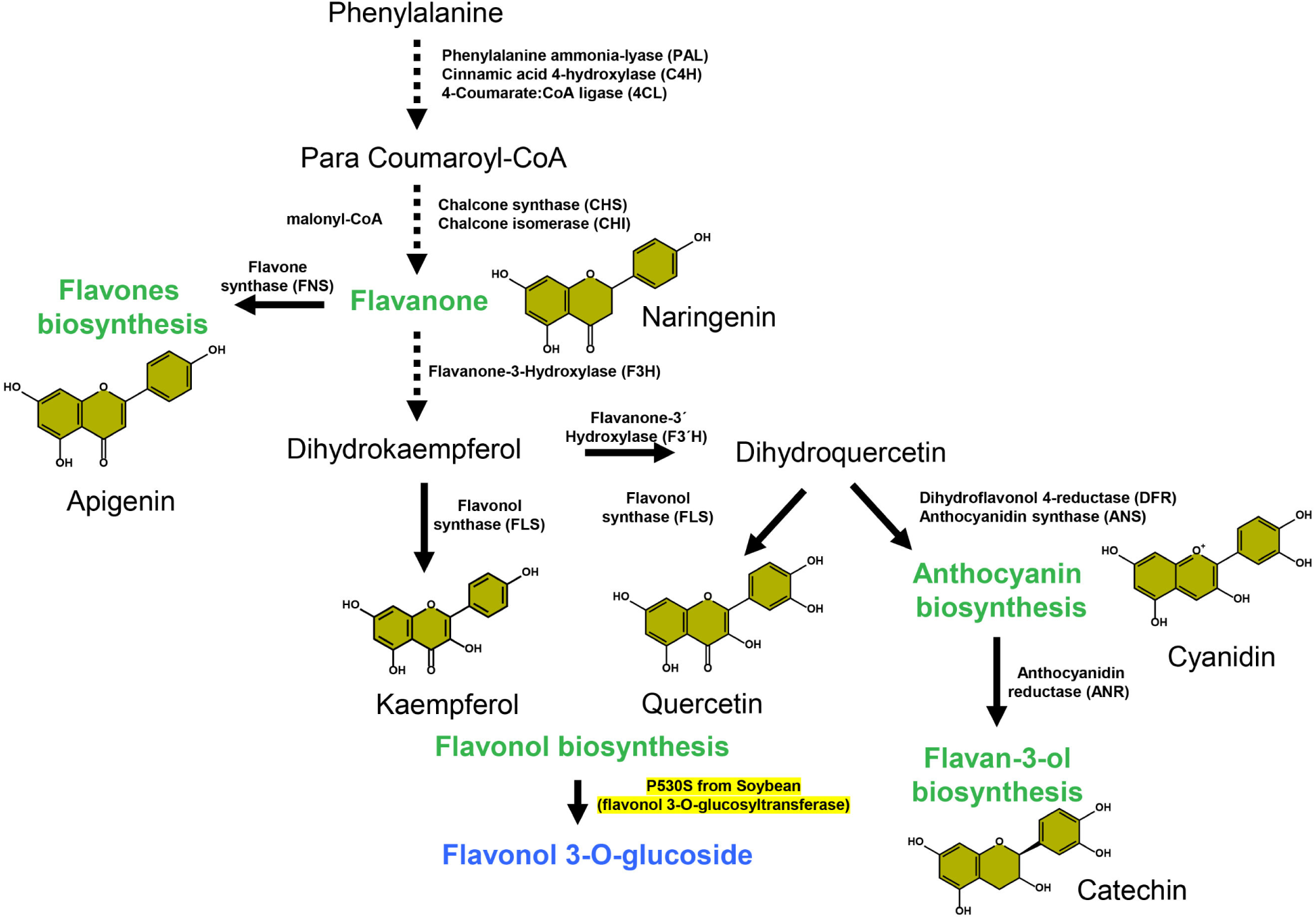
The proposed role of the *GmF3GlcT* protein product (highlighted in yellow) in the flavonoid biosynthetic pathway in higher plants. *GmF3GlcT* is flavonol-3-*O*-glucosyltransferase, with the greatest affinity for kaempferol, that forms flavonol 3-*O*-glucosides (in blue). *GmF3GlcT* also has an affinity for other flavonols, including quercetin. It does not use apigenin, naringenin, cyanidin or catechin as substrates.

## Discussion

Here, we report on the use of QTL mapping, near isoline development, and map-based cloning to identify the gene responsible for QTL-M resistance, and from there, the use of metabolomic and biochemical approaches to determine its mode of action and discern the nature of resistance.

Different approaches have been used to identify the gene underlying QTL M. Wang et al. (2015) used GWAS (Genome-Wide Association Study) to identify a calcium-responsive transcription factor as the candidate gene but never reported any validation of their candidate gene. Liu et al. (2016) used expression analysis to identify two jasmonic-acid-responsive genes (MYC2 and JAM2 homologs) that are induced by insect feeding as the candidate genes. They did not verify if these alleles are present in susceptible genotypes or confirm the candidates through knockouts or over-expression. We ruled out these candidate genes as they are present in some susceptible Recombinant Substitution Lines (RSLs), and because their gene sequences lack any SNPs uniquely present in resistant lines. Instead, our mapping, gene expression, and metabolomics analyses identify *Glyma07g14530* (*Glyma.07g110300* in v. 2 of the reference genome (W82v2)) as the causative gene.

QTL-M confers both antibiosis and antixenosis types of resistance (Rector et al., 2000b). Antibiosis means a plant causes detrimental effects on an insect. In antixenosis or non-preference resistance, plants discourage the presence of insects. QTL-M confers both antibiosis and antixenosis against a broad range of insects, suggesting that several metabolites could be involved. Once we identified *Glyma07g14530* as the causative gene, it became indisputable that flavonoids are involved in resistance.

Levels of some flavonoids and isoflavones increase in soybean leaves in response to defoliating insects (O’Neill et al., 2010), and flavonoids have long been suspected of playing a role in defoliator resistance in soybean, but conclusive proof has been elusive. Until now, it has never been clear if the different metabolites identified are the cause of resistance or simply due to genotypic differences between soybean varieties. For example, rutin is frequently implicated in resistance, but Jack, which is very susceptible to defoliating insects, has very high levels of rutin (data not shown). In this work, about 90% of the total flavonoid peak area from leaf extracts comprised flavonol conjugates, with kaempferol glucosides predominating in Benning and quercetin glucosides in Jack (**Fig 7A**).

Soybean leaves contain approximately 700 mg 100 g^-1^ dry weight flavonols (Romani et al., 2003) (Lee et al., 2022), consisting of their aglycones and their glycosylated products. Of these, kaempferol, quercetin, and isorhamnetin account for almost 80 to 90% of the total weight depending on the genotype of soybean. Glycosylation of flavonols mostly occurs at the 3rd position of the backbone with the addition of glucose, rhamnose, and galactose as the major modifications (Yang et al., 2018). The flavonols are decorated with two (50%) or three sugars (44%) to enhance their stability, solubility, and bioavailability, which further facilitates cellular transport, regulates biological activities such as plant development and stress response, and UV protection (Harborne and Williams, 2000, Mierziak et al., 2014). ‘Himeshirazu’ and Sodendaizu, both of which contain QTL-M, produce triglycosyls, specifically kaempferol 3-*O*-α-L-rhamnopyranosyl-(1→4)-[α-L-rhamnopyranosyl-(1→6)-β-D-galactopyranoside] and quercetin 3-*O*-β-D-glucopyranosyl-(1→2)-[α-L-rhamnopyranosyl-(1→6)-β-D-galactopyranoside] (Murai et al., 2013). The absence of triglucosyls but not other triglycosyls probably reflects the absence of a functional *F3GlcT* in QTL-M containing plants. In contrast, Zhao et al. (2015), looking at seed composition, attributed the increased resistance provided by QTL-M to elevated levels of genistein and glycitein.

Analyzed together, our data show that *Glyma07g14530* encodes a glucosyltransferase with a preference for flavonols with 3-hydroxy-2-phenyl-4H-chromen-4-one (3-Hydroxyflavone) as a minimal core structure. Plant UDP-glucosyltransferases belonging to the GT1 family of CAZymes (Drula et al., 2022) are known to catalyze the addition of glucose to the 3, 5, 7, 3′′ or 4′′ -OH positions. GlymaP530S does not have transferase activity with apigenin and luteolin, which lack the 3-OH group, but is active on a similar core-chemical structure with an additional 3-OH group, such as kaempferol, and to a lesser extent, quercetin, myricetin, and isorhamnetin, indicating that P530S is a selective flavonol 3-*O*-glucosyltransferase (**Fig 6**), coded by an ortholog of the *Arabidopsis atF3GlcT* or UG78D2 (Yin et al., 2012, Saito et al., 2013).

Complementing a QTL-M resistant line with *Glyma07g14530-S* restores susceptibility, and silencing *Glyma07g14530-S* results in resistance (**Fig 4**). Therefore, *Glyma07g14530* as present in most soybean germplasm is an allele for susceptibility. In wild-type soybean, insect feeding leads to upregulation of flavonol glucosylation, thus increasing the susceptibility of the host plant. The premature termination mutation found in PI 229358 prevents flavonol glucosylation, and thus is responsible for the leaf-chewing insect resistance associated with QTL-M.

Flavonoids are widely distributed in plants, and their core structure and bioactivity can be further modified by enzymes involved in modulation of hydroxylation, glycosylation, acylation, or methylation (Alseekh et al., 2020). Glycosylation by UGTs is an important regulator of flavonoid metabolism and accrual (Gachon et al., 2005). Of these, flavonols like kaempferol and quercetin with a C-ring 3-ketone/hydroxyl group and their glycosides affect both insect and stress resistance (Gautam et al., 2023, Simmonds, 2003, Agati et al., 2025).

‘IAC-17,’ another cultivar with QTL-M, has elevated levels of kaempferol-3-*O*-L-rhamnopyranosyl-glucopyranoside, rutin, and other flavanols. IAC-17 also has elevated levels of isorhamnetin, along with other flavonoids and proteinase inhibitors (Gómez et al., 2018). Although PA content was not measured, there is decreased gut proteolytic activity in caterpillars feeding on IAC-17, which is consistent with effects attributed to PA (Gómez et al., 2020).

For many metabolites like flavonols, glycosylation converts them from non-polar, insoluble molecules into a soluble glycosides that are actively recognized and sequestered into the vacuole for storage. As shown in **Fig 7**, in wild-type, insect-susceptible soybeans, dihydrokaempferol or dihydroquercetin are converted, respectively, to kaempferol or quercetin through a flavonol synthase (FLS). Next, a flavonol 3-*O*-glycosyltransferase (F3GT, specifically F3GlcT in this case) glucosylates its flavonol acceptor molecules, generating specific glucosides that are recognized by membrane transporters, allowing their storage in vacuoles. We postulate that PA-accumulating soybean mutants have a non-functional F3GT, leading to accumulation of flavonols. Specifically, FLS is sensitive to feedback inhibition (Yin et al., 2012), which could occur if a non-functional F3GT led to accumulation of flavone aglycones. In soybean, FLS inhibition should shunt metabolites into leucopelargonidin or leucocyanidin through the action of dihydrofolate reductase (DFR), leading to the eventual synthesis and accumulation of PAs, thus explaining why plants with QTL-M accumulate proanthocyanidins upon insect feeding. PAs, also known as condensed tannins, are known to both deter and be toxic to several insects (Harborne and Grayer, 1993).

In addition, the introgression of QTL-M leads to changes in the amounts of 378 metabolites upon insect feeding, and these changes might also contribute to resistance (Yousefi-Taemeh et al., 2021).

Finally, in field trials conducted under rainfed (i.e., non-irrigated) conditions, the line carrying QTL-M outyielded the best commercial checks by 33% (Mailhot, 2022). This result strongly suggests that the resistant allele of *Glyma07g14530* may confer a measure of drought tolerance in addition to insect resistance, which in turn may help explain the observed yield superiority of lines containing QTL-M, which is higher than would be expected from insect resistance alone, in regional field trials. Drought tolerance is frequently associated with flavonoids (Nayak et al., 2025). Protection from reactive oxygen species due to abiotic stress is associated with PA’s specifically and some other flavonoids (Gourlay et al., 2022). In the future, the direct modulation of flavonols through breeding to obtain drought and insect resistance should be a realistic goal.

## Methods

### Plant materials

The Japanese soybean landrace ‘Sodendaizu’ (PI 229358) is the source of the resistant allele of QTL-M. Benning is an elite soybean variety adapted to Georgia latitudes and that, as are almost all other soybean varieties, is susceptible to leaf-chewing insects. To generate the RSLs, a line of Benning^M^ (the superscript “M” indicates that the resistance-conferring version of QTL-M has been backcrossed into it.) was developed from Benning x PI 229358 through marker-assisted selection and then crossed to Benning to obtain a population of 1,991 BC_7_F_1:2_ plants. This population was screened as described by Zhu (2006), to identify heterozygous recombinant lines in the QTL-M region. These were self-pollinated to obtain homozygous Recombinant Substitution Lines (RSLs) (**Fig S1A**). Five of these were used to fine map the leaf chewing insect resistance locus. RSL 42, 47, 48, and 50 are resistant, and RSL 54 is susceptible.

The cultivar Jack and the BC_3_F_2_-derived Jack^M^ (Walker et al., 2004) were used for biolistic transformation. Jack, Jack^M^, Benning, and the BC_6_F_2:3_-derived Benning^M^ were used for the gene expression and proanthocyanidin assays.

### Preparation of insects for bioassays

SBL eggs were obtained from Benzon Research Inc. (Carlisle, PA). The eggs were incubated for 72 hr at 25C in a 600-mL (20 oz) clear polystyrene cup (Letica Corporation, Rochester Hills, MI, USA) sealed with a dome lid (Letica Corporation); each cup contained seven mL of plaster of Paris saturated with water to maintain 75% relative humidity. Neonate caterpillars were used to infest the bioassays.

### Identification of Ch.07 segment required for QTL-M resistance

Due to the fact we had no idea on what type of gene would lead to resistance, as well as the technology available when this work was started, we selected a map-based cloning approach.

The RSLs were screened for resistance in antixenosis assays with SBL, as described by Zhu et al. (2006) and genotyped for SNP and SSR loci between Sat_258 and Satt702, which define the QTL-M region (Zhu et al., 2006).

Polymorphic SNPs between Benning and PI 229358 within the QTL-M region were discovered using the soybean genome assembly version 4.0. Briefly, the 13,800-14,100 Mb segment from Chr.7 was used as a template to design 18 primer pairs that amplified 400-600-bp fragments every 50-100 kb (**Table S2**). DNA samples from PI 229358 and Benning were amplified with the 18 primer pairs in a 20-µL PCR reaction. Each reaction consisted of 2 µL of 40 ng µl ^-1^ genomic DNA, 1X PCR reaction buffer, 2.5 mM MgCl_2_, 100 µM of each dNTP, 0.2 µM of each primer, and 0.5 U of GoTaq Flexi DNA polymerase (Promega, Madison, WI). PCR reactions were performed in a T100^TM^ Thermal Cycler (Bio-Rad Laboratories Inc., Hercules, CA, USA). The PCR program consisted of an initial denaturation (94C for 5 min), followed by 35 cycles of 94° C for 30 sec, 58C for 30 sec, and 72C for 40 sec, and a final step of 72C for 5 min. The products were analyzed on a 1.5% agarose, 1X TBE gel to verify amplification of a single product. Each product was sequenced using the BigDye Terminator v3.1 Cycle Sequencing Kit (PE ABI, Foster City, CA). The amplicons were analyzed on an ABI 3730 automated sequencer (PE-ABI, Foster City, CA). For each fragment, the PI 229358 and Benning sequences were aligned to identify SNPs. Each RSL was then genotyped by amplifying and sequencing the SNP-containing fragments.

Sequences for the SSRs markers, Sat_425 and Satt729 (**Table S2**) were obtained from SoyBase (Grant et al., 2010) and were used to genotype the RSLs. For SSR genotyping, the PCR reactions were prepared using the protocol described by Li et al. (2002). The PCR products were analyzed on an ABI 3730 automated sequencer, and the data were processed with GeneScan v. 2.1 and Genotyper v. 2.5 software (PE ABI, Foster City, CA) using the procedures described by Li et al. (2002). The SNP and SSR marker data were used to build a graphical genotype of the QTL-M region for each RSL. The graphical genotypes were compared to their insect-resistance phenotypes to identify the PI 229358 introgression that is shared among all the resistant RSLs, and therefore contains QTL-M.

### PI 229358 BAC library screening, and BAC-clone sequencing

The PI 229358 BAC library reported by Zhu et al. (Zhu et al., 2009) was screened to identify BAC clones in the QTL-M region. High-density BAC filters were prepared as described by Zhu et al. (2009) and used in a hybridization-based screening of the library. Three DNA probes: two flanking, and one within the QTL-M region (**Fig S1C**), were used to screen the BAC filters. Primers designed from the Williams 82 reference genome (**Table S2**), were used to amplify the radiolabeled (^32^P) probes from PI 229358 genomic DNA. BAC-end sequences were obtained to determine the clone’s order in the QTL-M region contig based on the Williams 82 reference genome. Briefly, each clone was inoculated in 5 mL of LB broth supplemented with 12.5 µg mL^-1^ chloramphenicol and shaken at 280 rpm for 16 h at 37C. BAC DNA was isolated from the liquid cultures, using a standard alkaline lysis protocol. The BAC-DNA was used as template for sequencing with the universal primers T7 and M13 reverse, using BigDye Terminator v3.1 Cycle Sequencing Kit (PE ABI). Lastly, two overlapping clones covering QTL-M, (**Fig S1D**) were fully sequenced at the Clemson University Genomics Institute, and their assembled sequence was aligned to that of Williams 82 reference genome (**Fig S1E**).

### Annotation of BAC sequences

The sequences from the BAC clones 134P08 and 118D14 were assembled and annotated in Geneious version 8 (www.geneious.com). The primers S72_3610K-F and SNP13885-R, (**Table S2)** flanking the PI 229358 introgression required for QTL-M resistance (**Fig S1C**), were mapped to the BAC-clone sequence assembly. The PI 229358 sequence between S72_3610K-F and SNP13885-R was aligned to the corresponding sequence in Williams 82 (Glyma.Wm82.a1, Gmax1.01) using the mauveAligner algorithm (Darling et al., 2004). The soybean gene models (Wm82.a1.v1) were used to identify differences (SNPs and INDELs) in the gene-coding sequences that could be associated with resistance.

### Identification of candidate genes for QTL-M resistance

A panel of 34 insect-susceptible soybean accessions, including the 32 accessions that form most of the USA soybean ancestral germplasm pool, along with Benning and Jack, was used to narrow the number of candidate genes for resistance (**Fig 2SA**). Briefly, primers (**Table S2**) were designed to amplify the seven genes containing SNPs polymorphic between the susceptible Williams 82 and the resistant PI 229358 (**Table S1**). PCR products were amplified with KAPA HiFi polymerase (Kapa BioSystems, Boston, MA), using genomic DNA from each of the 32 insect-susceptible lines as template. PCR products were visualized in a 1X TAE + cytidine gel. Single products were extracted from the gel using the Zymoclean Gel DNA Recovery Kit (Zymo Research, Orange, CA). The purified PCR products were sequenced from both ends, using the gene-specific PCR primers. Sequencing reactions were carried out using the BigDye Terminator v3.1 Cycle Sequencing Kit (PE ABI) and analyzed on an ABI 3730 automated sequencer (PE-ABI). The sequences were quality-trimmed and mapped to each corresponding reference gene model in Geneious. Each sequence alignment was analyzed to determine the allele for each insect-susceptible accession at each SNP. All genes for which a SNP had the same allele in PI 229358 and any of the insect-susceptible accessions were excluded as candidates for insect-resistance.

A panel of 18 insect-resistant soybean accessions (Boethel, 1999, Kilen and Lambert, 1998), including cultivated and wild soybean, was used to further identify the candidate gene for QTL-M resistance (**Fig S2B**). Two gene models, *Glyma07g14470* and *Glyma07g14530* had SNP alleles unique to PI 229358 and were sequenced from insect-resistant accessions. The PCR, sequencing, and sequence analysis procedures were as described previously. Any gene sharing a SNP allele in common between PI 229358 and PI 227687 was further excluded as a candidate for insect-resistance, because PI 227687’s resistance does not map to QTL-M.

### DNA and RNA isolation

Genomic DNA was isolated from unexpanded trifoliolate leaves using the modified CTAB procedure described by Zhu et al. (2006). DNA samples were resuspended in 50 µl of TE/RNase buffer (10 mM Tris-HCL [pH 8.0], 1 mM EDTA, 100 µg mL^-1^ RNAse A). Leaf tissue harvested for RNA isolation was immediately frozen in liquid nitrogen. Total leaf RNA was isolated from 100 mg of tissue; the tissue was homogenized in Tri-Reagent Buffer (Sigma, St. Louis, MO), in a 2010 GenoGrinder (Spex SamplePrep, Metuchen, NJ). RNA samples were processed according to the manufacturer’s instructions. RNA samples were resuspended in 50 µL of RNAse-free water (Ambion, Austin, TX), and stored at −20 C.

### Expression of *Glyma07g14470* and *Glyma07g14530* in leaf-tissue

RNA isolated from Benning and Benning^M^ plants was used for RT-PCR to determine if the candidate genes are expressed in leaf-tissue. Briefly, five Benning^M^ and Benning plants were planted in 450-mL polystyrene cups using Fafard 2 mix (Conrad Fafard, Agawam, MA). Three holes were punched in the bottom of each cup for drainage. Plants were grown in an insecticide-free greenhouse under a 16-h/8-h light/dark photoperiod. To keep the plants in a vegetative stage, sunlight was supplemented with 400 J s^-1^ light provided by Phillips ED-18 high-pressure sodium bulbs (Phillips Inc., Andova, MA, USA). The temperature was maintained at approximately 28 C during the day and 20 C at night. Expanded trifoliolate leaves were collected for RNA isolation once the plants reached the V4 stage, and the plants were immediately infested with 30 neonate SBL caterpillars. Expanded insect-chewed trifoliolate leaves were collected for RNA isolation, 72 h after infestation. RNA was isolated as described previously, and the RNA extracts were treated with Turbo DNAse (Ambion) to remove contaminating DNA. RT-PCR primers for *Glyma07g14470* and *Glyma07g14530*, and the rest of the genes in QTL-M were designed in Geneious (**Table S2**). The metalloprotease gene, *Glyma03g29351*, was used as a constitutive gene control; RT-PCR primers are described by Libault (2008). RT-PCR primer sequences to amplify the wound-inducible pathogenesis-related 10 (PR10) gene were obtained from Graham (2003). First-strand cDNAs were synthesized in a 20-µL Superscript III Reverse Transcriptase (Invitrogen, Carlsbad, CA), containing 1 µg of RNA and 1 µL of 500 µg mL^-1^ Oligo(dT)_12-18_. First-strand cDNAs were used as templates in gene-specific PCR reactions. Each reaction consisted of 2 µL of first-strand cDNA, 1X PCR reaction buffer, 2.5 mM MgCl_2_, 100 µM of each dNTP, 0.2 µM of each primer, and 0.5 U of GoTaq Flexi DNA polymerase (Promega, Madison, WI). PCR reactions were performed in a T100^TM^ Thermal Cycler (Bio-Rad Laboratories Inc., Hercules, CA, USA), using a standard PCR program modified for the annealing temperature of each primer pair. PCR products were visualized in 1.5% TBE gels.

### Cloning of full-length cDNAs from *Glyma07g14470* and *Glyma07g14530*

To confirm the annotation of the *Glyma07g14470* and *Glyma07g14530* transcripts, rapid cDNA ends (RACE) PCR was performed using leaf RNA isolated from Benning and Benning^M^ as templates. The cDNAs were amplified using the SMARTer RACE cDNA kit (ClonTech, Mountain View, CA) according to the manufacturer’s instructions. The gene-specific primers described in **Table S2** were used for amplification and sequencing of the RACE products. Full-length transcripts corresponding to *Glyma07g14470* could not be amplified from leaf RNA samples due to the gene’s chimeric nature and homology with other soybean genes. As an alternative, *Glyma07g14470* was expressed in *Arabidopsis.* Briefly, the coding sequences of *Glyma07g14470* were amplified from Benning and Benning^M^ with KAPA HiFi polymerase (KAPA BioSystems) using the primers described in **Table S2**. The purified PCR products were cloned into the pDONR/Zeo plasmid vector (Invitrogen), by recombination between the *att*L1 and *att*L2 sites. The coding sequences were then transferred into the pEarleyGate202 expression vector by recombination between *att*B1 and *att*B2 sites. The *Arabidopsis* ecotype Landsberg erecta was transformed using*. tumefaciens* GV3101 carrying the pEarleyGate202 expression vectors. T_1_ transformants were selected with 50 mg L^-1^ BASTA (glufosinate ammonium). Leaf RNA was isolated from transformants as described previously. The expression of the transgenes was confirmed by RT-PCR as previously described. Rapid cDNA ends (RACE) PCR reactions were performed to clone the full-length transcripts. Finally, the 5’-RACE and 3’-RACE products obtained from *Glyma07g14470* in *Arabidopsis*, and *Glyma07g14530* in soybean, were assembled and compared to the corresponding annotated gene models in Geneious.

### qRT-PCR assays for *Glyma07g14530*

To determine if *Glyma07g14530* is induced by caterpillar damage, time-course qRT-PCR experiments were set up to measure *Glyma07g14530* transcript levels at 0, 24, 48, and 72 h after infestation. The first experiment was used to test whole plants. Briefly, four sets of five plants from each Benning, Benning^M^, Jack, and Jack^M^ were grown in a greenhouse as described previously. After the plants reached the V4 stage of development, 0-h leaf samples were collected for RNA isolation from the first set of plants from each genotype. Immediately after, the rest of the plants were infested with 30 SBL caterpillars each. Samples from sets 2, 3, and 4 of plants were collected at 24, 48, and 72 h after infestation, respectively. The second experiment was used to test detached leaves. Briefly, four sets of five plants from each Benning, Benning^M^, Jack, and Jack^M^ were grown in a greenhouse until they reached the V4 stage of development. Then, 0-h leaf samples were collected for RNA isolation from the first set of plants. Immediately after for the rest of the plants, a trifoliolate leaf was collected from each plant and placed in a 600-mL (20 oz) clear polystyrene cup (Letica Corporation, Rochester Hills, MI, USA) sealed with a dome lid (Letica Corporation); each cup contained plaster of Paris as described. Three neonate SBL caterpillars were added to each cup. Infested cups were placed in a growth chamber at 27C, and a 14-h light period was provided by fluorescent bulbs (T8 F032/730/Eco, Sylvania Octron, Danvers, MA, USA) that provided approximately 40 µmol photons m^-2^ s^-1^ (Zhu et al., 2008). Samples from sets 2, 3, and 4 of detached leaves, were collected at 24, 48, and 72 h after infestation, respectively.

Leaf RNA extractions were carried out as described previously. The RNA extracts were treated with Turbo DNase (Ambion) to remove contaminating DNA. Fifty ng of treated RNA were used as template for KAPA SYBR FAST One-Step qRT-PCR Kit (Kapa Biosystems). Briefly, 10-µl qRT-PCR reactions were performed in duplicate in a LightCycler 480II (Roche Diagnostics, Indianapolis, IN). The qRT-PCR program was 42C for 5 min and 95C for 3 min, for reverse transcription; and 45 amplification cycles of 95C for 10 sec, 65C for 20 sec, and 72C for 1 sec. The melting curve analysis, used to confirm the specificity of the qRT-PCR reactions, was set up from 60C to 95C, at a ramp rate of 0.11C second^-1^. The primer set 530qRT-F6 and 530qRT-R6 (**Table S2**) was used to amplify *Glyma07g14530*. The metalloprotease gene was used as a reference gene to normalize the expression of *Glyma07g14530*. The primers used to amplify the metalloprotease gene are the same as described previously. PCR efficiencies of each primer pair were determined using PCR reactions containing serial dilutions of each PCR product as template. Ct values for each qRT-PCR reaction were calculated using the Absolute Quantification analysis program. The ratio of target gene in comparison to the reference gene was calculated using the formula described by Pfaffl (2001).

### Vector construction and soybean transformation

Three plasmid vectors were used for complementation and silencing of *Glyma07g14530* in transgenic soybean lines (**Fig S3**). Two overexpression vectors used in the complementation assays were built as follows:

The coding sequences for *Glyma07g14530-S* (the wild-type, susceptible allele) and *Glyma07g14530***-**R (the resistant, mutant allele) were amplified from leaf DNA isolated from Benning and Benning^M^ respectively, with the primers *Asc*I-530-OE-F and *Avr*II-530-OE-R (**Table S2**) using the Phusion HF polymerase (New England Biolabs, Ipswich, MA). The pGmURNAi2 contains a gene-of-interest (GOI) expression cassette driven by the *G. max* ubiquitin (*GmUbi*) promoter (Hernandez-Garcia et al., 2009) and the *Pisum sativum* rubisco (rbcS) terminator. The GOIs were cloned between *GmUbi* and *rbcS* using the *Asc*I and *Avr*II restriction sites. After ligation, the plasmids were used to transform competent 10-beta *E. coli* cells (New England Biolabs); colonies were selected in solid LB medium supplemented with 50 µg mL^-1^ ampicillin. Positive colonies were confirmed by PCR and grown in liquid LB medium with ampicillin. Plasmids were purified with the GenElute Plasmid Miniprep (Sigma-Aldrich). The GmUbiP:Glyma530-R:rbcST, and GmUbiP:Glyma530-S:rbcST cassettes were verified by Sanger sequencing. A miRNA-induced gene silencing vector was built to silence *Glyma07g14530*. A 100-bp target fragment was amplified from *Glyma07g14530*’s coding sequence using the Phusion HF polymerase, with the primers *Asc*I-1510-530targ-F and *Avr*II-530targ-R (**Table S2**). The miR1510 target identified by Jacobs et al. (2016) was fused to the *Glyma07g14530* sequence in *Asc*I1510-530targ-F. The PCR product and the plasmid pGmURNAi2 were both digested with *Asc*I/*Avr*II. After ligation, the plasmids were used to transform competent 10-beta *E. coli* cells. Colony screening, miniprep, and sequencing reactions, to check the GmUbiP:1510:530:rbcST cassette were performed as described previously. For biolistic transformation of soybean, the expression and silencing cassettes were cloned into pSPH1 as I-*Ppo*I fragments. Plasmid SPH1 is derived from the pSMART HC Kan (Lucigen Corporation, Middleton, WI, AF532107), which was modified to contain the *hygromycin phosphotransferase* gene under the control of the *Solanum tuberosum* ubiquitin (StUbi3) promoter and terminator (StUbi3P:*hph*:StUbi3T), and the meganuclease I-*Ppo*I site.

To generate the transgenic soybean lines, somatic embryos from Jack and Jack^M^ were prepared and transformed using the procedure described by Trick et al. (Trick et al., 1997), with the following modifications. Somatic embryos were induced from immature cotyledons in Murashige and Skoog basal medium containing 40 mg L^-1^ 2,4-D. The resulting somatic embryos were transferred to half the concentration of 2,4-D for maintenance. Jack embryos were transformed with GmUbiP:Glyma530-R:rbcST and GmUbiP:1510:530:rbcST pSPH1 plasmids in separate events, to generate OE:530-R complementation lines and 1510:530 silencing lines respectively. Jack^M^ embryos were transformed with GmUbiP:Glyma530-S:rbcST and GmUbiP:1510:530:rbcST pSPH1 plasmids in separate events to generate OE:530-S complementation lines and 1510:530 silencing lines respectively.

For transformation, embryos were subjected to microprojectile bombardment, using 0.6-μm gold particles, with 10 ng of vector DNA per batch, shooting at the 7.58 MPa (1100 psi) setting. Briefly, one week after bombardment embryos were transferred to FNL medium containing 20 mg L^-1^ hygromycin for selection of transgenic tissue. Transgenic embryos were selected 6 to 8 weeks after bombardment and propagated in FNL medium. Once enough tissue was available, they were transferred to SHaM (soybean histodifferentiation and maturation) medium. Transgenic soybean plants (T_0_) were analyzed using standard molecular techniques to ensure the transgenes were present. For each transgenic line, the T_0_ plants were self-pollinated to obtain T_1_ seed. To determine the zygosity of T_1_ plants, the Invader assay (Hologic Inc., Madison, WI) (Gupta et al., 2008) was run on a Synergy 2 plate reader (BioTek Instruments Inc., Burlington, VA) according to manufacturer’s instructions. Homozygous T_1_ plants were self-pollinated to obtain T_2_ seeds. Selected T_2_ plants were self-pollinated to obtain T_3_ seeds.

### Characterization of transgenic soybean lines

Three independent lines each from Jack OE-530R, Jack 1510:530, Jack^M^ OE-530S, and Jack^M^ 1510:530 were characterized in bioassays measuring resistance against SBL caterpillars. The T_2_ generations were evaluated in greenhouse (choice, antixenosis) assays. The complementation lines, Jack OE-530R and Jack^M^ OE-530S, were evaluated in the first bioassay. Briefly, ten T_2_ plants from each line were grown as described previously. Leaf DNA was isolated from each plant, which was used as template for a PCR reaction with the primers Gmubi842-F and RbcSt110-R (**Table S2**) to confirm the presence of the transgenic cassette. Leaf RNA isolated from each PCR-positive plant was used in qRT-PCR reactions to measure the relative expression of the native *Glyma07g14530* and the transgenic construct OE:530S or OE:530R. The qRT-PCR reactions were set up as previously described. The primer set 530qRT-F6 and 530qRT-R6 was used to amplify the native *Glyma07g14530*, and the primer set qRbcSt-F and qRbcSt-R (**Table S2**) was used to amplify OE:530S or OE:530R constructs. Five T_2_ plants expressing both the native gene and transgene were used in the bioassay. The experimental design was a randomized complete block design with five replications. Each block contained one plant as the experimental unit. One plant from each complementation line, and one plant each of Jack and Jack^M^ were included in each block. The greenhouse bioassay was set up as described by Ortega et al. (2016a). Once the plants reached V4, each block was transferred to a 61 x 61 x 91 cm polyester-mesh cage (BioQuip products, Rancho Dominguez, CA, USA). Ten neonate SBL caterpillars were placed on each plant. Within a cage, caterpillars could move among the plants at will. The assay ended at ∼10 days, when defoliation of Jack reached 50%. The percent defoliation (Ortega et al., 2016a) of each plant was estimated by at least three persons, and the mean was used for an analysis of variance. The silencing lines Jack 1510:530 and Jack^M^ 1510:530 were evaluated in the second bioassay. Fifteen T_2_ plants from each line were grown and tested by PCR as described previously. Then, qRT-PCR reactions, set up as previously described, were used to measure the relative expression of the native *Glyma07g14530* and the transgenic construct 1510:530. To determine if the native gene was silenced in the transgenic lines expressing 1510:530, the expression level of *Glyma07g14530* on each line was compared to the average relative *Glyma07g14530* expression from five Jack or Jack^M^ plants. Five T_2_ plants in which the native *Glyma07g14530* was silenced were used in the bioassay, which was set up as described above for the complementation lines.

The T_3_ generations were evaluated in growth chamber (non-choice, antibiosis) assays. The complementation lines Jack OE-530R and Jack^M^ OE-530S were evaluated in the first bioassay. Ten T_3_ plants from each line were planted; the transgene’s presence and expression were determined by PCR and qRT-PCR, respectively, as described previously, and five plants per line were selected for the bioassay. The experimental design was a randomized complete block design with five replications. Each block contained one cup as the experimental unit. One cup from each complementation line, and one cup from each Jack and Jack^M^ were included in each block. The growth chamber bioassay was set up as described by Ortega et al. (2016a), once the plants reached V4 stage. Briefly, one trifoliolate leaf from each plant was placed into a corresponding 600-mL (20-oz) clear polystyrene cup (Letica Corporation) sealed with a dome lid and containing plaster of Paris as previously described. Each leaf was infested with five neonate SBL caterpillars. While feeding took place, the cups were placed in a growth chamber set at 27C. A 14-h light period was obtained using fluorescent bulbs (T8 F032/730/Eco, Sylvania Octron, Danvers, MA, USA) yielding ∼40 µmol photons m^-2^ s^-1^. The leaves in each cup were replaced with fresh leaves on the 4^th^ day, and afterwards, whenever 60% of the leaf area had been eaten. Percent defoliation was estimated based on the entire leaf (Ortega et al., 2016a). After seven days, caterpillars were immobilized at 4C for 24 h, after which they were weighed, and their mean weights were subjected to analysis of variance.

### Detection and profiling of abundant flavonoids

Ten milligrams of lyophilized leaf powder were sonicated at 4C in 500 µL methanol:chloroform (1:1, v/v) containing ^13^C_6_ trans-cinnamic acid and d_5_-benzoic acid as internal standards. Deionized water (200 μL) was added to the tubes for liquid-liquid phase partitioning, and the upper aqueous-methanol phase was used for analysis. Profiling was carried out by ultra-performance liquid chromatography coupled with quadrupole time-of-flight mass spectrometry (UPLC-QTOF). Extracts were filtered through a 0.2 μm PTFE (polytetrafluoroethylene) filter (Agilent Technologies) and resolved on an Agilent Zorbax Eclipse Plus C18 column (2.1 × 50 mm, 1.8 μm) using an Agilent 1290 Infinity II UPLC coupled to an Agilent 6546 LC/Q-TOF tandem MS. Mobile solvent A was water and 0.1% formic acid and mobile solvent B was acetonitrile with 0.1% formic acid. Flow rate was 0.5 mL min^−1^. The elution gradient was 3% B from 0–0.5 min, linear to 15% B over 3.0 min, to 30% B over 1.0 min, to 50% B over 1.0 min, to 100% B over 1.5 min with a 2-min hold followed by a 3-min post-column run. Column temperature was 45C. MS data were acquired in negative mode with gas temperature, 250C; nebulizer gas, 0.2759 MPa (40 psi); nozzle voltage, 500 V; and capillary voltage, 400 V. Exact mass data were processed using the MassHunter software suite Version 11.0 (Agilent). Compound IDs were obtained using the ID-browser function of MassProfiler v 10.0 (Agilent). The Agilent METLIN database (v B.08.00) was the primary library used for compound identification, supplemented with in-house authentic standards for further confirmation.

### Colorimetric assay for proanthocyanidins

To determine if QTL-M affects the levels of proanthocyanidins in soybean leaves, the Benning, Benning^M^, Jack, and Jack^M^ plants were analyzed using the butanol-HCl colorimetric assay described by Porter et al. (Porter et al., 1985). Briefly, 10 mg of ground freeze-dried tissue were transferred to a 2-mL Safe-Lock tube (Eppendorf, Hamburg, Germany). Next, 500 µL of methanol were added to each sample, and the samples were sonicated for 30 min in a FS-30 water-bath sonicator (Fisher Scientific, Pittsburg, PA). The tubes were spun in a tabletop centrifuge for 30 sec, and 400 µL of supernatant were transferred to a new tube. Six hundred µL of chloroform and 800 µL of water were added to each tube, in order. The tubes were vortexed and centrifuged for 1 min at 15000xg. The supernatant, containing the water-soluble proanthocyanidins, was transferred to a new tube. The samples were dehydrated in a Centrivap concentrator (Labconco Co., Kansas City, MO). Five hundred µL of butanol-HCl reagent (5% HCl in butanol) and 10 µL of 2% ferric reagent (2% ferric ammonium sulfate in 2M HCl) were added to each tube. The tubes were vortexed, and incubated at 95C for 20 min, and then cooled to room temperature. Each sample was read in duplicate wells, loading 100 µL per well, at 550 nm on a Synergy 2 plate reader (BioTek Instruments Inc).

### Expression and purification of recombinant GlymaP530S protein

To obtain GlymaP530S (the protein produced by the wild-type *Glyma07g14530* coding sequence), *Glyma07g14530* was cloned into pDEST-HisMBP (Plasmid #11085) obtained from Addgene using LR Clonase II from Thermo Fisher Scientific (USA) as per the manufacturer’s protocol and transformed into DH5α cells from Gene Choice Inc. (USA). The protein was expressed in *E. coli* Tuner (DE3) cells from Novagen (USA). Briefly, overnight cultures (5 mL) were used to inoculate 500 mL of Luria Broth (LB), and cultures were allowed to grow at 37C until OD_600_ reached ∼0.8. At this point, cultures were cooled on ice, and protein expression was induced by adding isopropyl β-d-thiogalactoside (IPTG; 200 µM) at 16C while shaking at 180 RPM for 18 hours. Cells were collected by centrifugation at 6370 × g for 10 min. The pellet obtained was resuspended resuspension buffer composed of 50 mM Tris-HCl, 400 mM NaCl, 1 mM BME, and 1% (v/v) glycerol and lysed using Avastin Emulsiflex C3 (Canada). The protein was purified from the clarified lysate using an MBPTrap HP column (Cytiva; USA) using the AKTA Pure 25 M chromatography system (Cytiva; USA). The protein fractions were concentrated to 10 mL with a 10 kDa molecular cut-off membrane from an Amicon Ultra centrifugal filter (Millipore, USA). Size exclusion chromatography was carried out using a HiLoad 16/600 Superdex 200 pg column (Cytiva; USA) using 50 mM Tris-HCl, 400 mM NaCl, 1 mM BME, and 1% glycerol. The purified protein was concentrated to 1 mg mL^-1^, based on an extinction coefficient of 121085 molar per centimeter, and stored at 4C for immediate use or at −80C for long-term storage. The purity of the purified protein was confirmed by sodium dodecyl sulfate-polyacrylamide gel electrophoresis (SDS-PAGE).

### UDP-Glo glycosyltransferase assay

Possible acceptors were screened using the UDP-Glo^TM^ assay kit (Promega, USA) to measure the production of the co-product UDP in reactions containing UDP-Glc as a donor and a series of pooled acceptor substrate groups to increase throughput for reaction optimization. The highest activity was determined among two acceptor groups consisting of apigenin, kaempferol, myricetin, isorhamnetin, and daidzein (Acceptor 21 to 24); genistein, gallocatechin, and epigallocatechin (Acceptor 25 to 28) (**Fig S6a**). Next, acceptors from groups 21 to 24 and 25 to 28 were assayed individually, leading to the identification of kaempferol as having the highest activity (**Fig S6b)**. From this point forward, kaempferol was used in subsequent reactions to determine donor specificity, pH, and temperature optima. Gly-NaOH with pH 8 showed the highest glucosyltransferase activity (**Fig S7a**), and 35.8C was the optimum temperature (**Fig S7b**). Using the strategy of (Sheikh et al., 2017), different UDP-sugars were screened without an acceptor substrate. Of the seven UDP-sugars tested, UDP-Glc showed the highest hydrolytic activity (**Fig 6**). Finally, using optimized conditions, all the available acceptors were screened for glucosyltransferase activity; the highest activity was determined using kaempferol as the acceptor substrate.

Reactions (30 µL) were carried out at 30C for an hour and consisted of UDP-glucose (100 µM), DTT (1 mM), and 6 µg of purified P530S protein in the indicated buffer. UDP-sugar specificity was evaluated by setting up assays as described with the indicated UDP-sugars (100 µM) in Tris-HCl (100 mM, pH 7.5), DTT (1 mM), and kaempferol (50 µM). The pH optimum was evaluated by incubating UDP-glucose (100 µM), DTT (1 mM), and kaempferol (50 µM) in various buffers, with the pH varying from 5.8 to 11. Finally, using the optimized conditions obtained, all the acceptors were evaluated by glo-assay. The UDP byproduct of GT reactions was quantified using the UDP-Glo^TM^ assay kit according to the manufacturer’s instructions using 384-well Corning 4513 white plates with luminescence measured using a Synergy LX Multi-mode microplate reader (BioTek, USA). The released UDP was quantified using a standard curve.

### Matrix-assisted laser desorption ionization time-of-flight (MALDI-TOF) mass spectrometry (MS) analysis

Matrix-Assisted Laser Desorption/Ionization Time-of-Flight (MALDI-TOF) mass spectra of P530S enzyme reaction products were acquired using a Microflex LT spectrometer (Bruker, USA). Reactions (40 µL) were carried out for 16 h at 35.8C using 24 µg of the P530S enzyme, UDP-Glc (1 mM), and kaempferol (0.125 mM), DTT (1 mM) in glycine-NaOH (100 mM, pH 8.0). For MS of reaction products, 1 µL of the reaction was mixed with 1 µL matrix (2.5 mg of norharman in 80% acetonitrile) directly on the plate and air-dried. Negative-ion spectra from 1000 laser shots were averaged to generate the spectrum for each sample.

### Statistical analysis

Data recorded from qRT-PCRs, bioassays, HPLC analysis, and the condensed tannin assay were analyzed using JMP version 10.0 (SAS Institute, Inc., Cary, NC). The normality of each dataset was tested with the Shapiro-Wilk test (*P* < 0.05). The data were then subjected to a one-way ANOVA (*P* < 0.05) followed by a post-hoc Tukey-Kramer multiple comparison test (*P* <0.01) to determine significant differences between treatments.

## Supporting information

Loss of flavonol 3-O-glucosyltransferase activity confers soybean resistance to 2 leaf-chewing insects - Supplemental Figures & Tables

## Author contributions

P.K.P. made the constructs for protein expression, optimized expression, purified protein, performed biochemical characterization, and drafted the manuscript. M.A.O. sequenced the candidate genes in the insect-resistant accessions, cloned *Glyma07g14530*, characterized the transgenic soybean lines, designed and performed all the bioassays, and helped draft the manuscript. B.K.H. developed SNPs to genotype the QTL-M region in the RSLs, screened the BAC library and sequenced BAC clones, identified candidate genes based on Williams 82 sequence, and sequenced the candidate genes in the susceptible panel. H.R.B developed the near isolines to make this study possible. P.R.L. designed and built the plasmid vectors for soybean transformation. C.J.T. and S.A.H. assisted with the molecular screens, the bioassays, and the overall interpretation of results. B.R.U. generated the initial protein expression constructs, provided overall guidance, funding and edited the manuscript. W.A.P. conceived and directed the overall study and helped draft and edit the manuscript.

## Acknowledgements

We thank Donna Tucker for her assistance in soybean tissue culture and generation of transgenic events, Alexander Sterling Graf and Dylon Jacob Quiros for assisting in protein expression and purification. Thanks also to Eudald Illa-Berenguer for assisting with the graphs.

## Supporting information

**Fig 1 QTL-M region in soybean chromosome 7.**

**Fig 2. Identification of candidate genes for QTL-M insect resistance.**

**Fig 3. Development of transgenic soybean lines,**

**Fig.4. Weight gain in soybean loopers caterpillars feeding on overexpression or silencing lines**

**Fig 5. SDS PAGE of purified P530S.**

**Fig 6. Acceptor substrate screen for P530S.**

**Fig 7. pH (a) and temperature (b) screen of P530S activity**

**Table 1. Polymorphisms between Williams 82 and PI 229358 for gene models contained in the QTL-M region.**

**Table 2. List of primer sets used**

---

**Fig 7.**
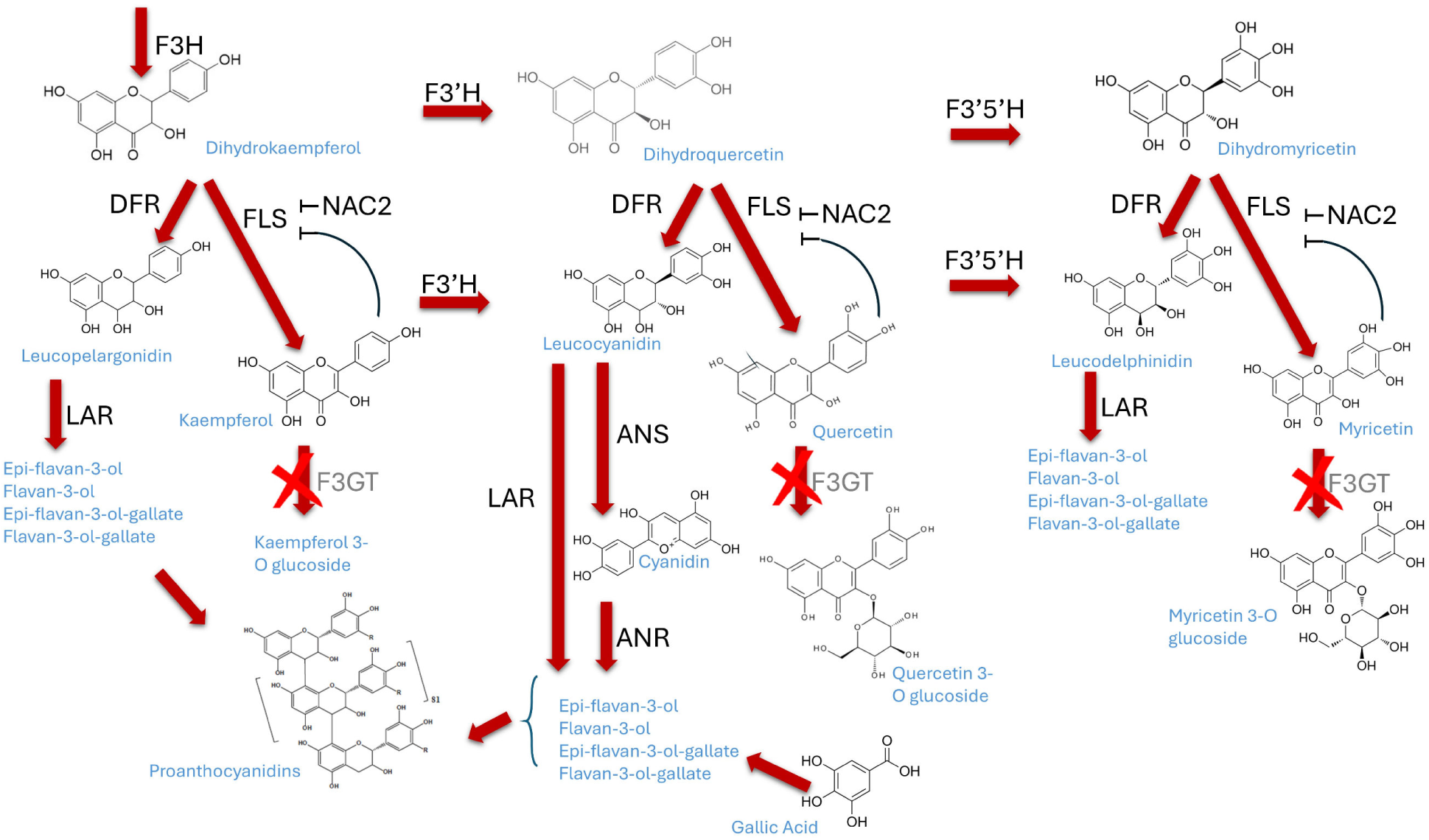
The proposed mechanism for proanthocyanidin formation upon loss of flavonol-3-*O*-glucosyltransferase activity. In wild-type soybean, dihydrokaempferol or dihydroquercetin are converted, respectively, to kaempferol or quercetin through a flavonol synthase (FLS). A flavonol 3-O-glycosyltransferase (generically called F3GT) glucosylates the flavonols, allowing their storage in a vacuole. FLS is regulated by the NAC2 transcription factor, and is sensitive to feedback inhibition. In soybean, FLS inhibition would shunt metabolites into leucopelargonidin or leucocyanidin through the action of dihydrofolate reductase (DFR), leading to the eventual synthesis and accumulation of proanthocyanins (condensed tannins). Drawing based on Saito et al. (2013), Yin et al. (2012), and Yu et al. (2023).

## References

Agati, G., Brunetti, C., Nascimento, L.B.D., Gori, A., Lo Piccolo, E. & Tattini, M. 2025. Antioxidants by nature: an ancient feature at the heart of flavonoids’ multifunctionality. New Phytologist, 245, 11–26.

Alseekh, S., De Souza, L.P., Benina, M. & Fernie, A.R. 2020. The style and substance of plant flavonoid decoration; towards defining both structure and function. Phytochemistry, 174.

Boethel, D.J. 1999. Assessment of soybean germplasm for multiple insect resistance, CRC Press, Boca Raton, FL.

Crop-Protection-Network Estimates of crop yield losses due to diseases and invertebrate pests: an online tool. 2024 ed.: Crop Protection Network.

Darling, A.C.E., Mau, B., Blattner, F.R. & Perna, N.T. 2004. Mauve: multiple alignment of conserved genomic sequence with rearrangements. Genome Research, 14, 1394–403.

Deutsch, C.A., Tewksbury, J.J., Tigchelaar, M., Battisti, D.S., Merrill, S.C., Huey, R.B. & Naylor, R.L. 2018. Increase in crop losses to insect pests in a warming climate. Science, 361, 916–919.

Dillon, F.M., Chludil, H.D. & Zavala, J.A. 2017. Solar UV-B radiation modulates chemical defenses against larvae in leaves of field-grown soybean. Phytochemistry, 141, 27–36.

Drula, E., Garron, M.L., Dogan, S., Lombard, V., Henrissat, B. & Terrapon, N. 2022. The carbohydrate-active enzyme database: functions and literature. Nucleic Acids Research, 50, D571–D577.

Eulgem, T. & Somssich, I.E. 2007. Networks of WRKY transcription factors in defense signaling. Current Opinion in Plant Biology, 10, 366–371.

Gachon, C.M.M., Langlois-Meurinne, M. & Saindrenan, P. 2005. Plant secondary metabolism glycosyltransferases: the emerging functional analysis. Trends in Plant Science, 10, 542–549.

Gautam, H., Sharma, A. & Trivedi, P.K. 2023. The role of flavonols in insect resistance and stress response. Current Opinion in Plant Biology, 73.

Gillen, A.M. 2021. Table 127 - General summary of performance preliminary test VII 2020 *Uniform Soybean Tests Southern States*. USDA-Agricultural Research Service.

Gillen, A.M. 2023.Table 114 - General summary of performance uniform test VII 2022. Uniform Soybean Tests Southern States. Stoneville, Mississippi 38776: USDA-Agricultural Research Service.

Gómez, J.D., Pinheiro, V.J.M., Silva, J.C., Romero, J., Meriño-Cabrera, Y., Coutinho, F.S., Lourençao, A.L., Serrao, J.E., Vital, C.E., Fontes, E.P.B., Oliveira, M.G.A. & Ramos, H.J.O. 2020. Leaf metabolic profiles of two soybean genotypes differentially affect the survival and the digestibility of caterpillars. Plant Physiology and Biochemistry, 155, 196–212.

Gómez, J.D., Vital, C.E., Oliveira, M.G. & Ramos, H.J. 2018. Broad range flavonoid profiling by LC/MS of soybean genotypes contrasting for resistance to *Anticarsia gemmatalis* (Lepidoptera: Noctuidae). PloS one, 13, e0205010.

Gourlay, G., Hawkins, B.J., Albert, A., Schnitzler, J.P. & Constabel, C.P. 2022. Condensed tannins as antioxidants that protect poplar against oxidative stress from drought and UV-B. Plant Cell and Environment, 45, 362–377.

Graham, M.Y., Weidner, J., Wheeler, K., Pelow, M.J. & Graham, T.L. 2003. Induced expression of pathogenesis-related protein genes in soybean by wounding and the *Phytophthora sojae* cell wall glucan elicitor. Physiological and Molecular Plant Pathology, 63, 141–149.

Grant, D., Nelson, R.T., Cannon, S.B. & Shoemaker, R.C. 2010. SoyBase, the USDA-ARS soybean genetics and genomics database. Nucleic Acids Research, 38, 843–846.

Gupta, M., Nirunsuksiri, W., Schulenberg, G., Hartl, T., Novak, S., Bryan, J., Vanopdorp, N., Bing, J. & Thompson, S. 2008. A non-PCR-based Invader® assay quantitatively detects single-copy genes in complex plant genomes. Molecular Breeding, 21, 173–181.

Harborne, J.B. & Grayer, R.J. 1993. Flavonoids and insects. In: Harborne, J. B. (ed.) The Flavonoids : advances in research since 1986. 1st ed. London; New York: Chapman & Hall.

Harborne, J.B. & Williams, C.A. 2000. Advances in flavonoid research since 1992. Phytochemistry, 55, 481–504.

Hernandez-Garcia, C.M., Martinelli, A.P., Bouchard, R.A. & Finer, J.J. 2009. A soybean (*Glycine max*) polyubiquitin promoter gives strong constitutive expression in transgenic soybean. Plant Cell Reports, 28, 837–849.

Jacobs, T.B., Lawler, N.J., Lafayette, P.R., Vodkin, L.O. & Parrott, W.A. 2016. Simple gene silencing using the trans-acting siRNA pathway. Plant Biotechnology Journal, 14, 117–127.

Kilen, T.C. & Lambert, L. 1998. Genetic control of insect resistance in soybean germplasm PI 417061. Crop Science, 38, 652.

Lee, S., Kim, H.W., Lee, S.J., Ha Kwon, R., Na, H., Kim, J.H., Choi, Y.M., Yoon, H., Kim, Y.S., Wee, C.D., Yoo, S.M. & Lee, S.H. 2022. Comprehensive characterization of flavonoid derivatives in young leaves of core-collected soybean (*Glycine max* L.) cultivars based on high-resolution mass spectrometry. Scientific Reports, 12.

Li, Z., Wilson, R.F., Rayford, W.E. & Boerma, H.R. 2002. Molecular mapping genes conditioning reduced palmitic acid content in N87-2122-4 soybean. Crop Science, 42, 373–378.

Libault, M., Thibivilliers, S., Bilgin, D.D., Radwan, O., Benitez, M., Clough, S.J. & Stacey, G. 2008. Identification of four soybean reference genes for gene expression normalization. The Plant Genome, 1, 44.

Liu, H.L., Che, Z.J., Zeng, X.R., Zhang, G.Z., Wang, H. & Yu, D.Y. 2016. Identification of single nucleotide polymorphisms in soybean associated with resistance to common cutworm (*Spodoptera litura* Fabricius). Euphytica, 209, 49–62.

Mailhot, D.J. 2022. Yield summary: MG IV-VIII soybean variety performance. Georgia Statewide Variety Testing.

Mierziak, J., Kostyn, K. & Kulma, A. 2014. Flavonoids as important molecules of plant interactions with the environment. Molecules, 19, 16240–16265.

Murai, Y., Takahashi, R., Rodas, F.R., Kitajima, J. & Iwashina, T. 2013. New flavonol triglycosides from the leaves of soybean cultivars. Natural Product Communications, 8, 1934578X1300800410.

Narvel, J.M., Walker, D.R., Rector, B.G., All, J.N., Parrott, W.A. & Boerma, H.R. 2001. A retrospective DNA marker assessment of the development of insect resistant soybean. Crop Science, 41, 1931.

Nayak, J.K., Mohanty, D., Mahapatra, D., Mondal, S., Mohanty, A., Sangeeta, S., Nayak, G., Senapaty, J. & Panda, B. 2025. Flavonoids: a natural shield of plants under drought stress. In: Aires, A. (ed.) Plant Secondary Metabolites - Occurrence, Structure and Role. IntechOpen.

O’neill, B.F., Zangerl, A.R., Dermody, O., Bilgin, D.D., Casteel, C.L., Zavala, J.A., Delucia, E.H. & Berenbaum, M.R. 2010. Impact of elevated levels of atmospheric CO_2_ and herbivory on flavonoids of soybean (*Glycine max* Linnaeus). Journal of Chemical Ecology, 36, 35–45.

Oerke, E.C. 2006. Crop losses to pests. Journal of Agricultural Science, 144, 31–43.

Ortega, M.A. 2016. Cloning and using a QTL for insect resistance in soybean. PhD, University of Georgia.

Ortega, M.A., All, J.N., Boerma, H.R. & Parrott, W.A. 2016a. Pyramids of QTLs enhance host-plant resistance and Bt-mediated resistance to leaf-chewing insects in soybean. Theoretical and Applied Genetics, 129, 703–715.

Ortega, M.A., Davis, A.J., Boerma, H.R. & Parrott, W.A. 2016b. Suitability of soybean meal from insect-resistant soybeans for broiler chickens. Journal of Agricultural and Food Chemistry, 64, 2209–2213.

Ortega, M.A., Lail, L.A., Wood, E.D., All, J.N., Li, Z.L., Boerma, H.R. & Parrott, W.A. 2017. Registration of two soybean germplasm lines containing leaf-chewing insect resistance QTLs from PI 229358 and PI 227687 introgressed into ‘Benning’. Journal of Plant Registrations, 11, 185–191.

Pagano, M.C. & Miransari, M. 2016. The importance of soybean production worldwide. Abiotic and Biotic Stresses in Soybean Production. Amsterdam; Boston: Elsevier/AP, Academic Press is an imprint of Elsevier.

Pfaffl, M.W. 2001. A new mathematical model for relative quantification in real-time RT-PCR. Nucleic Acids Research, 29, e45–e45.

Porter, L.J., Hrstich, L.N. & Chan, B.G. 1985. The conversion of procyanidins and prodelphinidins to cyanidin and delphinidin. Phytochemistry, 25, 223–230.

Rector, B.G., All, J.N., Parrott, W.A. & Boerma, H.R. 2000a. Quantitative trait loci for antibiosis resistance to corn earworm in soybean. Crop Science, 40, 233–238.

Rector, B.G., All, J.N., Parrott, W.A. & Boerma, H.R. 2000b. Quantitative trait loci for antixenosis resistance to corn earworm in soybean. Crop Science, 40, 531–538.

Ribeiro, A.V., Führ, F.M., Menger, J.P., Havill, J.S. & Koch, R.L. 2025. Legume host range and soybean host plant resistance for soybean tentiform leafminer, *Macrosaccus morrisella* (Lepidoptera: Gracillariidae). Journal of Economic Entomology, 118, 1731–1741.

Romani, A., Vignolini, P., Galardi, C., Aroldi, C., Vazzana, C. & Heimler, D. 2003. Polyphenolic content in different plant parts of soy cultivars grown under natural conditions. Journal of Agricultural and Food Chemistry, 51, 5301–5306.

Saito, K., Yonekura-Sakakibara, K., Nakabayashi, R., Higashi, Y., Yamazaki, M., Tohge, T. & Fernie, A.R. 2013. The flavonoid biosynthetic pathway in *Arabidopsis*: Structural and genetic diversity. Plant Physiology and Biochemistry, 72, 21–34.

Sheikh, M.O., Halmo, S.M., Patel, S., Middleton, D., Takeuchi, H., Schafer, C.M., West, C.M., Haltiwanger, R.S., Avci, F.Y., Moremen, K.W. & Wells, L. 2017. Rapid screening of sugar-nucleotide donor specificities of putative glycosyltransferases. Glycobiology, 27, 206–212.

Simmonds, M.S. 2003. Flavonoid-insect interactions: recent advances in our knowledge. Phytochemistry, 64, 21–30.

Skendzic, S., Zovko, M., Zivkovic, I.P., Lesic, V. & Lemic, D. 2021. The impact of climate change on agricultural insect pests. Insects, 12.

Smith, C.M. & Clement, S.L. 2012. Molecular bases of plant resistance to arthropods. Annual Review of Entomology*, Vol* 57, 57, 309–328.

Trapero, A., Ahrazem, O., Rubio-Moraga, A., Jimeno, M.L., Gómez, M.D. & Gómez-Gómez, L. 2012. Characterization of a glucosyltransferase enzyme involved in the formation of kaempferol and quercetin sophorosides in *Crocus sativus*. Plant Physiology, 159, 1335–1354.

Trick, H.N., Dinkins, R.D., Santarém, E.R., Samoyolov, R.D.V., Meurer, C., Walker, D., Parrott, W.A., Finer, J.J. & Collins, G.B. 1997. Recent advances in soybean transformation. Plant Tissue Culture and Biotechnology, 3, 9–26.

Van Duyn, J.W., Turnipseed, S.G. & Maxwell, J.D. 1971. Resistance in soybeans to the Mexican bean beetle. I. Sources of resistance. Crop Science, 11, 572–573.

Walker, D.R., Narvel, J.M., Boerma, H.R., All, J.N. & Parrott, W.A. 2004. A QTL that enhances and broadens Bt insect resistance in soybean. Theoretical and Applied Genetics, 109, 1051–1057.

Wang, H., Yan, H.L., Du, H.P., Chao, M.N., Gao, Z.J. & Yu, D.Y. 2015. Mapping quantitative trait loci associated with soybean resistance to common cutworm and soybean compensatory growth after defoliation using SNP marker-based genome-wide association analysis. Molecular Breeding, 35, 1–15.

Yang, B., Liu, H.L., Yang, J.L., Gupta, V.K. & Jiang, Y.M. 2018. New insights on bioactivities and biosynthesis of flavonoid glycosides. Trends in Food Science & Technology, 79, 116–124.

Yin, R., Messner, B., Faus-Kessler, T., Hoffmann, T., Schwab, W., Hajirezaei, M.R., Von Saint Paul, V., Heller, W. & Schäffner, A.R. 2012. Feedback inhibition of the general phenylpropanoid and flavonol biosynthetic pathways upon a compromised flavonol-3--glycosylation. Journal of Experimental Botany, 63, 2465–2478.

Yonekura-Sakakibara, K., Higashi, Y. & Nakabayashi, R. 2019. The origin and evolution of plant flavonoid metabolism. Frontiers in Plant Science, 10.

Yousefi-Taemeh, M., Lin, J., Ifa, D.R., Parrott, W. & Kovinich, N. 2021. Metabolomics differences of *Glycine max* QTLs resistant to soybean looper. Metabolites, 11.

Yu, K., Song, Y., Lin, J. & Dixon, R.A. 2023. The complexities of proanthocyanidin biosynthesis and its regulation in plants. Plant Commun, 4, 100498.

Zhang, Y., Guo, W., Chen, L., Shen, X., Yang, H., Fang, Y., Ouyang, W., Mai, S., Chen, H., Chen, S., Hao, Q., Yuan, S., Zhang, C., Huang, Y., Shan, Z., Yang, Z., Qiu, D., Zhou, X., Cao, D., Li, X. & Jiao, Y. 2022. CRISPR/Cas9-mediated targeted mutagenesis of GmUGT enhanced soybean resistance against leaf-chewing insects through flavonoids biosynthesis. Frontiers Plant Science, 13, 802716.

Zhao, G., Jiang, Z., Li, D., Han, Y., Hu, H., Wu, L., Wang, Y., Gao, Y., Teng, W., Li, Y., Zeng, G., Meng, F. & Li, W. 2015. Molecular loci associated with seed isoflavone content may underlie resistance to soybean pod borer (*Leguminivora glycinivorella*). Plant Breeding, n/a-n/a.

Zhu, S., Saski, C.A., Boerma, H.R., Tomkins, J.P., All, J.N. & Parrott, W.A. 2009. Construction of a BAC library for a defoliating insect-resistant soybean and identification of candidate clones using a novel approach. Plant Molecular Biology Reporter, 27, 229–235.

Zhu, S., Walker, D.R., Boerma, H.R., All, J.N. & Parrott, W.A. 2006. Fine mapping of a major insect resistance QTL in soybean and its interaction with minor resistance QTLs. Crop Science, 46, 1094.

Zhu, S., Walker, D.R., Boerma, H.R., All, J.N. & Parrott, W.A. 2008. Effects of defoliating insect resistance QTLs and a *cry1Ac* transgene in soybean near-isogenic lines. Theoretical and Applied Genetics, 116, 455–63.

