## Supplementary material for "Loss of flavonol 3-*O*-glucosyltransferase activity confers soybean resistance to leaf-chewing insects": Loss of flavonol 3-O-glucosyltransferase activity confers soybean resistance to 2 leaf-chewing insects - Supplemental Figures & Tables

### Supporting Information

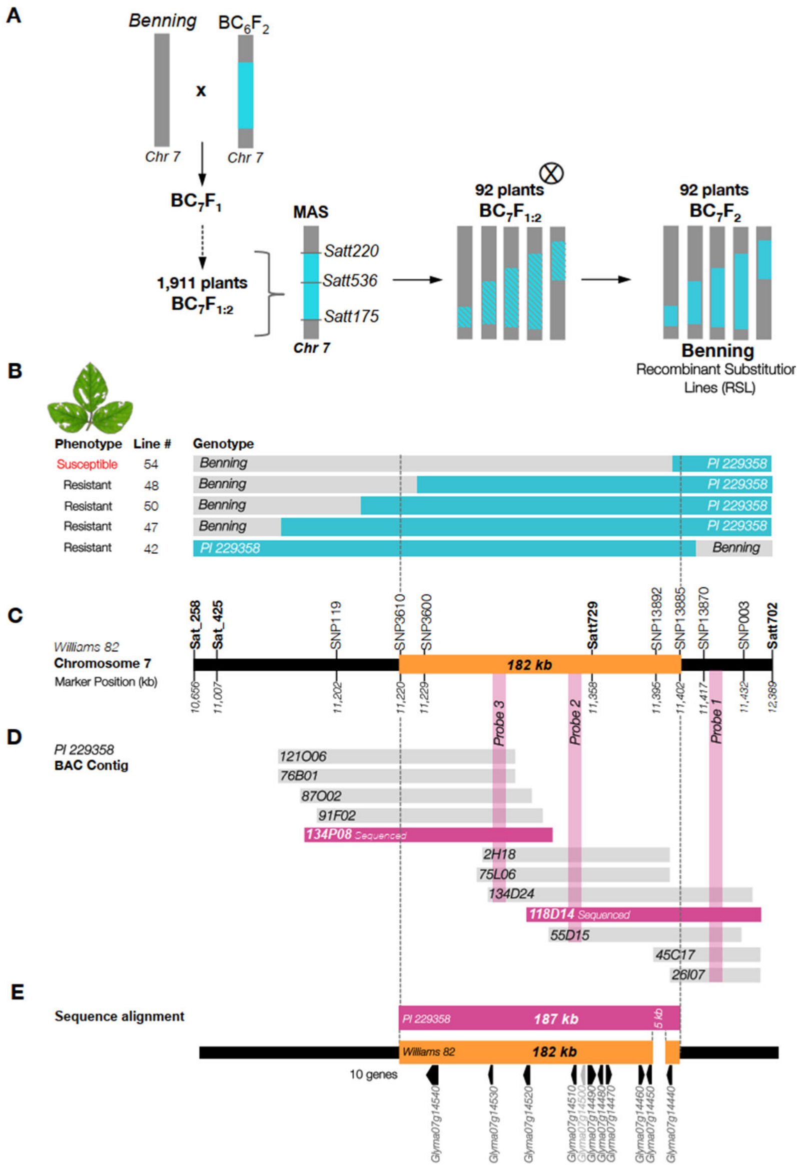


**Supporting Fig 1** **QTL-M region in soybean chromosome 7.** (**A**) Development of RSLs derived from Benning x PI 229358 for positional cloning of QTL-M. (**B**) Graphical genotypes of five RSLs. Grey and blue indicate loci carrying Benning and PI 229358 alleles, respectively. (**C**) Chr.7 segment containing QTL-M. The bar shows the position of molecular markers in Williams 82. All resistant lines contain the PI 229358 introgression between SNP3610 and SNP13885. (**D**) PI 229358 BAC-clone contig. Twelve clones were identified by hybridization with DNA probes; the contig was assembled using BAC-end sequences. Full-length sequences were obtained from two overlapping clones. (**E**) PI 229358 sequence aligned to Williams 82. Arrows indicate ten gene models and a pseudogene annotated in this region.

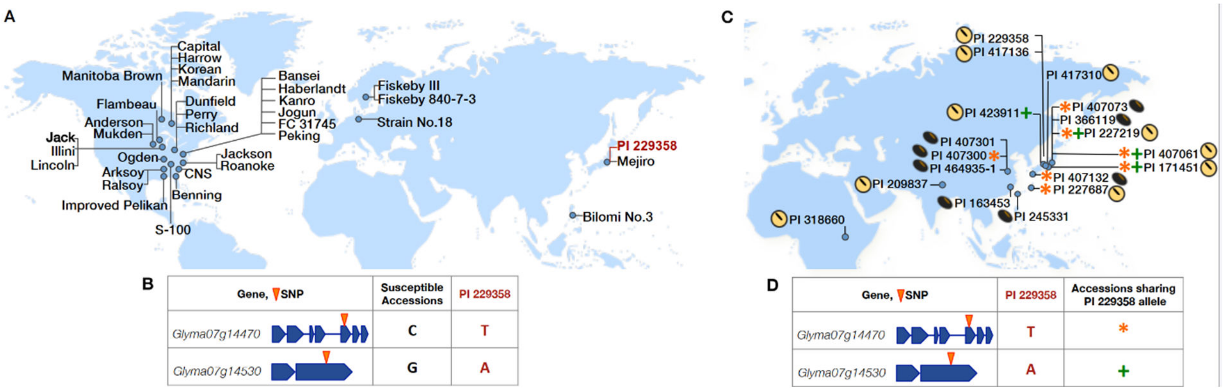


**Supporting Fig 2. Identification of candidate genes for QTL-M insect resistance.** (**A**) Panel of insect-susceptible soybeans. Polymorphic genes between PI 229358 and Williams 82 were sequenced in each accession to identify SNP loci carrying alleles unique to PI 229358. (**B**) Two gene models were identified that contained containing SNPs unique to PI 229358. (**C**) Panel of insect-resistant soybeans. Yellow seeds and brown seeds indicate cultivated and *G. soja* accessions respectively. (**D**) Accessions carrying the PI 229358 allele for *Glyma07g14470* and *Glyma07g14530* SNPs are indicated by an asterisk and a cross, respectively. These two SNPs are also found in other insect-resistant genotypes.

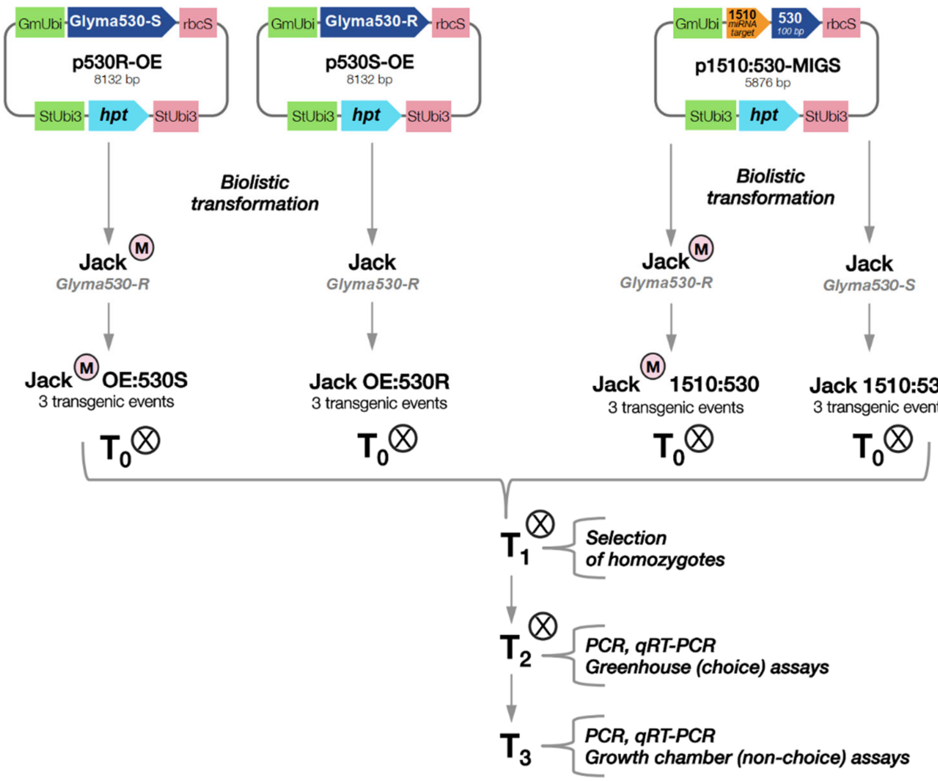


**Supporting Fig 3. Development of transgenic soybean lines.** The over-expression vectors p530S-OE and p530R-OE were used to generate complementation lines for Jack$^M$ and Jack, respectively. The silencing vector p1510:530-MIGS was used to generate silencing lines for both Jack$^M$ and Jack. $T_0$ and selected $T_1$ plants were self-pollinated. $T_2$ and $T_3$ plants were characterized for resistance against SBL caterpillars.

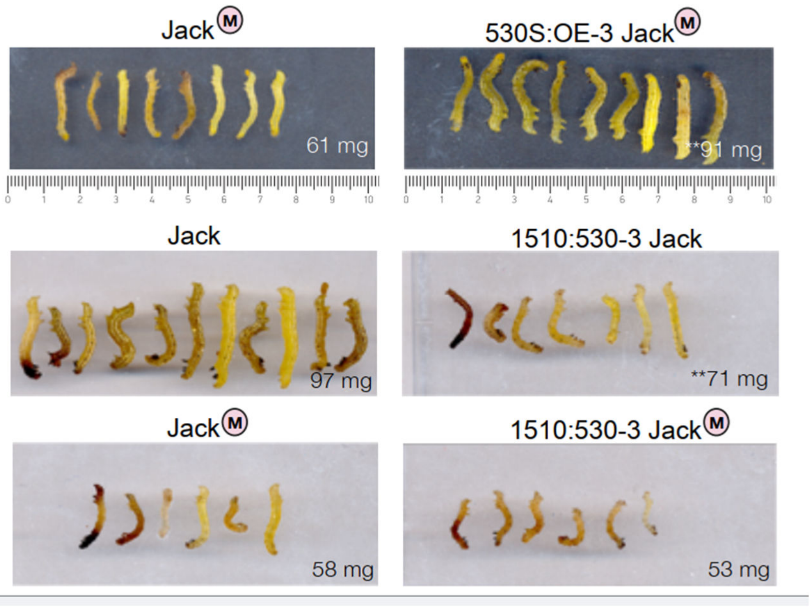


**Supporting Fig.4. Weight gain in soybean loopers caterpillars feeding on overexpression or silencing lines.** Caterpillars are significantly larger after seven days of feeding on 530S:OE-3 Jack$^{M}$ compared to Jack$^{M}$ (top row). SBL caterpillars are significantly smaller after feeding on 1510:530-3 Jack for 7 days compared to Jack (middle row). There is no difference in size of SBL feeding on Jack$^{M}$ and 150:530-3 Jack$^{M}$ $T_3$ lines (bottom row). 530S:OE-3 Jack$^{M}$ overexpresses the allele for susceptibility, while 150:530-3 Jack$^{M}$ has its allele for susceptibility silenced.

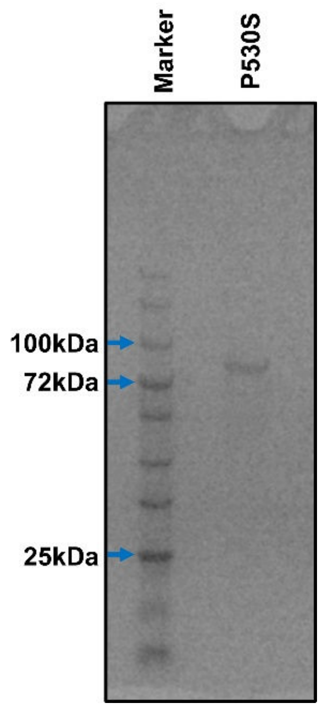


**Supporting Fig 5.** **SDS PAGE of purified P530S.** Five µg of purified protein were loaded onto a 12% SDS-PAGE gel. The purified protein migrates at the expected weight of 97.5 kD.

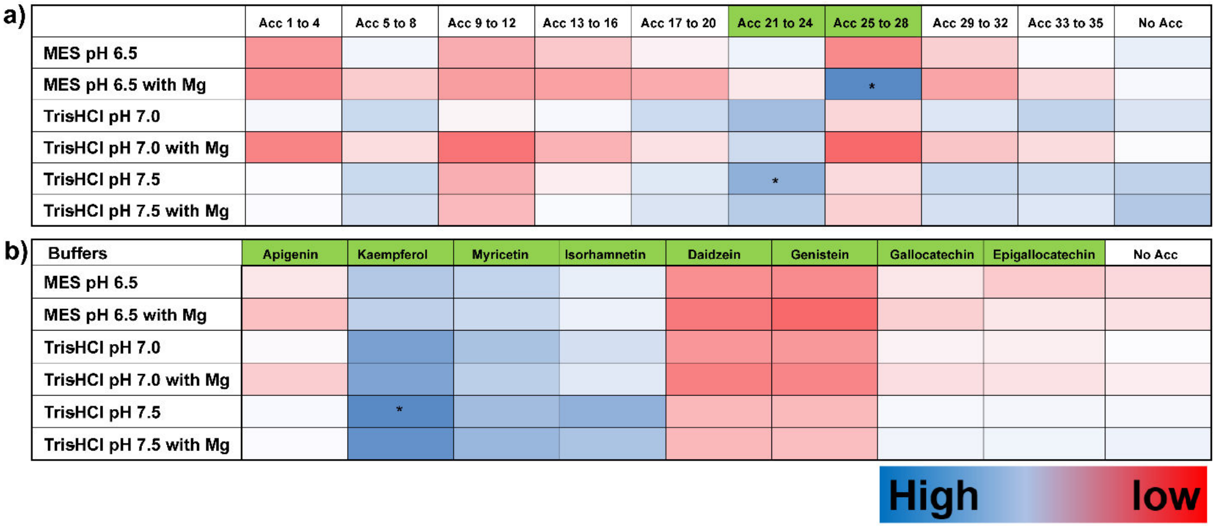


**Supporting Fig 6.** **Acceptor substrate screen for P530S** (a) Heat map showing the acceptor substrate screen for P530S glycosyltransferase activity using UDP-Glc as donor and pooled acceptor (Acc) substrates with indicated reaction conditions: Acc 1 to 4 (Eriodyctiol, Sativoside, 2-Hydroxynaringenin, & DL-Dihydrozeatin), Acc 5 to 8 (DL-4-Hydroxymandelonitrile, Benzyl alcohol, Mandelonitrile & Benzoic Acid), Acc 9 to 12 (Phloretin, Myrtillin, Geraniol, & Cyanidin), Acc 13 to 16 (Steviolbioside, Cis-Zeatin, Zeatin & Trans-Zeatin), Acc 17 to 20 (Eriodyctiol, Luteolin, Quercetin, & Naringenin), Acc 21 to 24 (Apigenin, Kaempferol, Myricetin, & Isorhamnetin), Acc 25 to 28 (Daidzein, Genistein, Gallocatechin, & Epigallocatechin), Acc 29 to 32 (Catechin, Epicatechin, Epigallocatechin Gallate, & Hydroquinone), Acc 33-35 (3,4 Dichloroaniline, 2,4,5 Trichlorophenol, 4-Chlorothiophenol). No acceptor or Hydrolysis is represented as No acc and Control (No P530S) is subtracted from each value. * Represents the substrate(s) with the highest activity on the heat map and will be used for future experiments. (b) Heat map to define substrate selectivity for Glyma530S using UDP-Glc as donor and acceptor substrates 21-28 as indicated. The final concentration of buffer used is 100 mM and mM magnesium in each case.

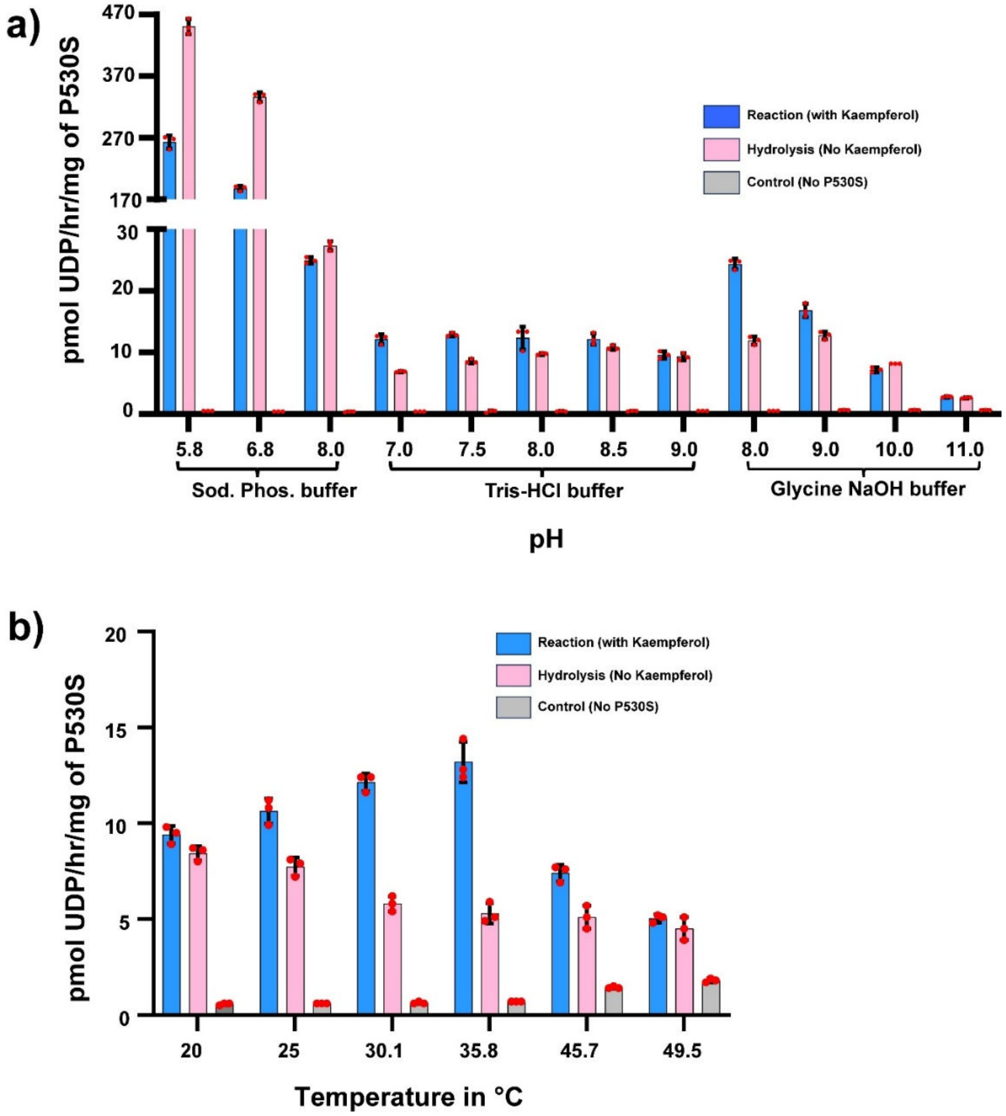


**Supporting Fig 7.** **pH (a) and temperature (b) screen of P530S activity**. Kaempferol is used as an acceptor, and UDP-Glc is used as a donor. Control represents the background without P530S (the protein from the allele that confers susceptibility). Activity is represented as the mean ± standard deviation for n =3 technical replicates (red circles).

**Supporting Table 1. Polymorphisms between Williams 82 and PI 229358 for gene models contained in the QTL-M region.**

| Gene ID | Predicted annotation[1] | Size | Williams 82/PI 229358 polymorphisms[3] |
|---|---|---|---|
| Glyma07g14440 | Unknown function | 1,220 bp | $^{832}$T/C ($^{278}$stop/R) |
| Glyma07g14450 | Unknown function | 1,424 bp | Same sequence |
| Glyma07g14460 | Cytochrome P450 | 2,610 bp | T/A, T/-, $^{726}$A/C ($^{242}$silent) |
| Glyma07g14470 | Ploop-NTPase | 1,903 bp | $^{500}$G/A ($^{167}$P/L) |
| Glyma07g14480 | MYB-related | 2,014 bp | $^{78}$G/C ($^{26}$K/N), G/A, A/T, A/C, $^{639}$T/A ($^{213}$silent) |
| Glyma07g14490 | Phosphoglycerate mutase | 3,128 bp | C/A, A/T, G/A |
| Glyma07g14500 | Pseudogene, Helix-loop-helix | 378 bp | Same sequence |
| Glyma07g14510 | Glucosyl/Glucuronosyl transferase | 1,705 bp | Same sequence |
| Glyma07g14520 | Unknown function | 1,891 bp | A/G |
| Glyma07g14530 | Glucosyl/Glucuronosyl transferase | 1,401 bp | $^{747}$G/A ($^{249}$W/stop) |
| Glyma07g14540 | DNAJ/HSP40 | 3,703 bp | Same sequence |

[1] Annotations based on the Wm82.a1.v1 gene models (www.soybase.org).

[2]Blue arrows represent exons, and blue lines indicate introns. Polymorphisms are indicated by orange triangles; details for each polymorphism are shown in the next column.

[3]For SNPs and/or INDELs in introns, the Williams 82 allele is shown first, followed by the PI 229358 allele. For SNPs in exons, the SNP position (bp) in the coding sequence is shown first, followed by the Williams 82 and PI 229358 allele; the effect of each polymorphism in the amino acid sequence is shown in parentheses.

**Supporting Table 2. List of primer sets used**

| PCR products to identify SNPs in the QTL-M region | Primer name | Sequence (5'->3') |
|---|---|---|
| S72_3420K | S72_3420K-F | CCTAACTCTCTTTAACCTCGTC |
| | S72_3420K-R | CTTGGTTGGGAAGTTCTG |
| S72_3430K | S72_3430K-F | AGGCTATAGATTAAGAATGCTAAGTC |
| | S72_3430K-R | CTTCACATTTCCTCTTATTAACAACC |
| S72_3450K | S72_3450K-F | TGATGTGATGTGATGTGACG |
| | S72_3450K-R | CCTCTATGTATTTCAGAAATGTCTC |
| S72_3490K | S72_3490K-F | CAAATGTCAATGCAATAATAGCGT |
| | S72_3490K-R | GAAGGTTGATAGTAATGACTGGAG |
| S72_3510K | S72_3510K-F | AAGGTGTATCCAAGATTAGCC |
| | S72_3510K-R | TCTGATTAGAAAGTCAATGATTCCC |
| S72_3530K | S72_3530K-F | GATTGTCCAACGTTCAGG |
| | S72_3530K-R | ACTTGTCTAACACATTGATGCTA |
| S72_3550K | S72_3550K-F | TTTGGTTATGTGTGGCTGG |
| | S72_3550K-R | CTTCTAGTAGTTAAGGTCTTCCC |
| S72_3570K | S72_3570K-F | GCTTCAACTTCTCTTCCTATCC |
| | S72_3570K-R | GTAGGGCATAGAGACGCA |
| S72_3600K | S72_3600K-F | ACACCAGCACAAAGTCC |
| | S72_3600K-R | GTCTCATTCAGTTCTCGCA |
| S72_3410K | S72_3410K-F | TTGTGCAGATATGAAGCTCTTTAG |
| | S72_3410K-R | CCACCTTCACAATCACG |
| S72_3422K | S72_3422K-F | CAGGTGCTCTGTGTTGT |
| | S72_3422K-R | CTGGTTCACTGACCTTCG |
| S72_3430K | S72_3430K-F | CCAAGTCAGGTATGAGAATGC |
| | S72_3430K-R | TCCACATCCAACACAATCG |
| S72_3443K | S72_3443K-F | CTCCATAGTTCAAGTACAACCC |
| | S72_3443K-R | ACCATCCATCTTCATGTACG |
| S72_3452K | S72_3452K-F | TTTGCTACATGGCAGCTT |
| | S72_3452K-R | GACACTGAAACACAATGCC |
| S72_3589K | S72_3589K-F | CTGGATGATTGTGAGGAGAC |
| | S72_3589K-R | AGAACTCGTTGAATCGCT AT |
| S72_3600K | S72_3600K-F | ACAGAACATTGCATTCCCTA |
| | S72_3600K-R | GCTGATATGTTCACAAAGCCA |
| S72_3610K | S72_3610K-F | CAGGGACATGATATGATTAGCAA |
| | S72_3610K-R | GGTCACACCTTCCATATCTC |
| S72_3616K | S72_3616K-F | CCATACAATGTTGCACACG |

| | | |
|---|---|---|
| | S72_3616K-R | CATATTAGTATACCAGGTGACGG |
| SNP003 | SNP003-F<br>SNP003-F | CATACTGATGATGGGCCG<br>GCCACTTCTCACACGTTT |
| Satt729 | Satt729-F<br>Satt729-R | AAGTGCTAGCCACAAGGTTGA<br>GATTTTGGTTTGCCCTTAGC |
| SNP119 | SNP119-F<br>SNP119-R | ATACAGCTCAATAGGTAGTACCAAT<br>TGGGAAAGGGTGCTTGATTCTA |
| SNP13870 | SNP13870-F<br>SNP13870-R | TGCCGATAGTGTCAATGAGT<br>ATTAATGTGATTGTTTGGACCTG |
| SNP13885 | SNP13885-F<br>SNP13885-R | GCTACTTGAACACAATCGATG<br>ACCTAACTTATTAGAATTTGCGTGA |
| SNP13892 | SNP13892-F<br>SNP13892-R | TAACTACTTGTATCACACTAGCATCT<br>ACTAAGTGTCTTGGATCGGA |
| SNP116 | SNP116-F<br>SNP116-R | CAGACATAAGTCCTCGATCAC<br>CCTAACGACAAAGGCCC |
| Sat_425 | Sat_425-F<br>Sat_425-R | CCTTTGAGACGCAACTGAAAAT<br>TGATGGTGGTGTTGGTGTAA |
| **Sequencing primers for candidate SNPs** | | |
| Glyma07g14460 | P450-SEQ-F1<br>P450-SEQ-R1<br>P450-SEQ-F2<br>P450-SEQ-R2<br>P450-SEQ-F3<br>P450-SEQ-R3<br>P450-SEQ-F4<br>P450-SEQ-R4<br>P450-SEQ-F5<br>P450-SEQ-R5<br>P450-SEQ-F6<br>P450-SEQ-R6<br>P450-SEQ-F7<br>P450-SEQ-R7 | AGCCATATCCATCCAAAGAC<br>TGGACCTTTGAGGAAACG<br>AGCTCATCTCAGCCTTCATA<br>GGAAGGACAACACCACTAA<br>GTGGTGGAATCTCACGTT<br>TTGGATGAGGCTGCTTAGT<br>TCAATTCAATGCTTTGGTGGAT<br>CATAAAGAAACAATGAACTGCCG<br>TCATCTTAGTGTAGTGTGATTCTTAGT<br>TGCAGCAATGAGAAGCC<br>TGAAATCTTTGCAAGCATCATAAC<br>CAATCGATCTCTGGAAAAGG<br>AATATGGACTCATCTGCTTAGG<br>AACTCTTGGAAAGGTGCTAAA |
| Glyma07g14470 | PLNTP-SEQ-F1<br>PLNTP-SEQ-R1<br>PLNTP-SEQ-F2<br>PLNTP-SEQ-R2<br>PLNTP-SEQ-F3<br>PLNTP-SEQ-R3<br>PLNTP-SEQ-F4 | ACACACATTCTAAGA CGGTTAC<br>GTCGTTGGAGGAGAGGT<br>GGATCAGTTACCACACCG<br>ACAATGAGCACCTGCAAATA<br>CCATTCTTCTACGGATTAAATAGTGT<br>ATGCACCTGTTTCTTGAGAT<br>TGGACTTCAGGGAGTTAGGA |

| | | |
|---|---|---|
| | PLNTP-SEQ-R4 | TCAAGTAGTGTTAATTAGTTGTTGGG |
| | PLNTP-SEQ-F5 | TGGACTTCAGGGAGTTAGGA |
| | PLNTP-SEQ-R5 | TCAAGTAGTGTTAATTAGTTGTTGGG |
| | PLNTP-SEQ-F6 | CAGAAGTATTGCTAGTTGTGG |
| | PLNTP-SEQ-R6 | CACAACACATCACAGAAAGAC |
| Glyma07g14480 | MYB-SEQ-F1 | GGGTAATTTCCACATTATTAATGGTT |
| | MYB-SEQ-R1 | AACGAAGACGACACGAC |
| | MYB-SEQ-F2 | AGAAGAAGATGAGATGCTACTGA |
| | MYB-SEQ-R2 | ACACAAATGCAAGTAACAACCATAAA |
| | MYB-SEQ-F3 | TGCTTCACATTGATCCTGG |
| | MYB-SEQ-R3 | CAAATTGTGCCTGCAACTC |
| | MYB-SEQ-F4 | CTAACACTGCAAGGGACG |
| | MYB-SEQ-R4 | GCTATGTCTTGTTTAAGGAAATTTGTA |
| | MYB-SEQ-F5 | CAGCAACTCCTAAATCGC |
| | MYB-SEQ-R5 | CCTCAGAATTATCAATGTAGGGC |
| | MYB-SEQ-F6 | CCCAGAGAATGTAGTAGTAGAC |
| | MYB-SEQ-R6 | CCACAGGGAAATCATCGAA |
| | MYB-SEQ-F7 | TGCATTTCCTCAGATTCCAG |
| | MYB-SEQ-R7 | GAATATATCATTGCAGAATGCGT |
| | MYB-SEQ-F7-1 | TTGCCCACAATTGATGAGC |
| | MYB-SEQ-R7-1 | TTGCAGAATGCGTTGATACC |
| | MYB-UP-SEQ-F1 | CTCTTTCTCCAACCCACG |
| | MYB-UP-SEQ-R1 | CAAGTCCTCCTTCCTCCCTAA |
| | MYB-UP-SEQ-F2 | GTTTGGAGGAAGAGTTGGT |
| | MYB-UP-SEQ-R2 | ACATATAAATAGTAAATAAGTCTCCCAGG |
| | MYB-UP-SEQ-F3 | TGATGTTGTCTTTTGTATTTGTCTTC |
| | MYB-UP-SEQ-R3 | TTATTCACTTGCATTCTCCACC |
| | MYB-UP-SEQ-F4 | GCCATGTTACCTGTACACT |
| | MYB-UP-SEQ-R4 | TTTACCACCCAACTTCATTATACT |
| | MYB-UP-SEQ-F5 | AGGGATTCCACAAAATATCTATACAAA |
| | MYB-UP-SEQ-R5 | ACGGTTATTCATGGCAGT |

| | | |
|---|---|---|
| Glyma07g14500 | HLH-SEQ-F1<br>HLH-SEQ-R1<br>HLH-SEQ-F2<br>HLH-SEQ-R2<br>HLH-SEQ-F3<br>HLH-SEQ-R3<br>HLH-SEQ-F4<br>HLH-SEQ-R4<br>HLH-SEQ-F5<br>HLH-SEQ-R5<br>HLH-SEQ-F6<br>HLH-SEQ-R6 | GTCACAACATCTTTAATTGATTAGGT<br>AAAGATCAATAACATCAAGAGTGC<br>CTACAGTTCATCAATCTGCGA<br>GCATATTGCACAGGTAGAGT<br>CATGGGATAAGAGGAACGC<br>CTAAATAGATGTACTGCTCGCT<br>AGAAAGGCACTCCATTTACT<br>CTATCTCCAGCCATTGTTATCG<br>AGTTTGAGTGAGTGAGGG<br>TAAGCATCTCAGCCACG<br>TGTGAGAAGAAAGGTTGCG<br>TGGATGATTAAGCGAAACCGA |
| Glyma07g14510 | UDP1-SEQ-F1<br>UDP1-SEQ-R1<br>UDP1-SEQ-F2<br>UDP1-SEQ-R2<br>UDP1-SEQ-F3<br>UDP1-SEQ-R3<br>UDP1-SEQ-F4<br>UDP1-SEQ-R4<br>UDP1-SEQ-F5<br>UDP1-SEQ-R5<br>UDP1-SEQ-F6<br>UDP1-SEQ-R6 | ATGAATTTCCACAGACCGA<br>GGATTAGTGGTAGAGAACGAG<br>CAATTCTTGAGTTCTCTAAGCG<br>CTTGTATGCTACACCTGATCG<br>ATATTTCCCATCCACGGC<br>GCATATACAGAAGGGATTCCTC<br>AATCTTTGGGTCCACTACG<br>TGAGCAAGGATTTGAACCTG<br>AGATTCTTGTGGGTGTTAAGAC<br>TCCACTTTAATGCCAACTGT<br>TGATGGTTTGAAAGTGGCTC<br>ACAACAGACATATTATGCAAGTG |
| Glyma07g14530 | UDP2-SEQ-F1<br>UDP2-SEQ-R1<br>UDP2-SEQ-F2<br>UDP2-SEQ-R2<br>UDP2-SEQ-F3<br>UDP2-SEQ-R3<br>UDP2-SEQ-F4<br>UDP2-SEQ-R4 | ATGCAGCAGTACCTTTGAC<br>ACAAGGGAAGTAGATGTAGGATA<br>CTAAACAACAATGGAGTCTCTG<br>CCAAATGACACATAAAGAACTGA<br>AGCTACCCTCCTGTGTAT<br>CTAGTGTCAACGTTTGGTCT<br>CTGAACAAAGAACCAATGCG<br>ATACACAATAGTTGACTTTATCTCATACA |
| **Endpoint RT-PCR primers** | | |
| Glyma07g14440 | Glyma07g14440-F1 | TTTCGTTTCTTTTGGAATCGCC |
| | Glyma07g14440-R1 | GTTGCAAATGTAAGGCTCCTC |
| Glyma07g14450 | Glyma07g14450-F1 | CAGGGCCATAAAAAGATATTTCATGGGGTC |
| | Glyma07g14450-R1 | GCTTCCAACAGCCCAAAAGACGCATAAAC |

| | | |
|---|---|---|
| Glyma07g14460 | Glyma07g144<br>60-F1 | CTGATTTGCTTGCACTTTTGCG |
| | Glyma07g144<br>60-R1 | CTTAGGGCCAACAAGCTCAAGGG |
| Glyma07g14470 | pLoop-NTP-F2 | CCCAGCAACTAGCATGCAACTTTTTCC |
| | pLoop-NTP-R2 | TATTGGCTGGACTTCAGGGAGTT |
| Glyma07g14480 | Glyma07g144<br>80-F3 | TCTCTTGCCCACAATTGATGAGC |
| | Glyma07g144<br>80-R3 | CTGAAAGAAAGGGAACCCAATGG |
| Glyma07g14490 | Glyma07g144<br>90-F1 | TCAAAGCCCCAATTTCCATCTT |
| | Glyma07g144<br>90-R1 | GAGGTGGTTCTTGCAGTGT |
| Glyma07g14500 | Glyma07g145<br>00-F4 | AAGCATTCCAACTTGGGCTTGAAGGAA |
| | Glyma07g145<br>00-R4 | GGTGGACCAAAGATGAACGTGGCTGAGATG |
| Glyma07g14510 | Glyma07g145<br>10-F3 | TTCTATAAACCATGTCCACATTATGTGATC |
| | Glyma07g145<br>10-R3 | CCCCCAAGCTTAAGTTTTTCCAC |
| Glyma07g14520 | Glyma07g145<br>20-F2 | GGCTGATGAAAAGGTGCATCAAT |
| | Glyma07g145<br>20-R2 | TCTTTGCCTTCTCCCTCCAC |
| Glyma07g14530 | Glyma07g145<br>30-F3 | GTGAGTATAGAGATCACCCAAAC |
| | Glyma07g145<br>30-R3 | CTTCCAATTCCATGAAGCTATTG |
| Glyma07g14540 | Glyma07g145<br>40-F1 | GCATGTTGAGAAGGGTATGC |
| | Glyma07g145<br>40-R1 | GATGGCTCATCATCATCTTCG |
| **Full-length cDNA cloning** | | |
| Glyma07g14470 | Glyma07g144<br>70-R3 | AAGTTGCATGCTAGTTGCTGGG |
| | Glyma07g144<br>70-R5 | CTGTCTTGTCAATTACAATCCA |
| | Glyma07g144<br>70-F10 | GCAAACAAACCTCTTTGACACTC |
| | Glyma07g144<br>70-F11 | CACATATGAGAAGGTCACATG |

| | | |
|---|---|---|
| | Glyma07g14470-R13 | GACCGTCGTGGGTTGGAGAAAGAGTTCCTC |
| Glyma07g14530 | 530-5-RACE | GGAGACGTTTTGCGAACTCGAGGATTG |
| | 530-3-RACE | GCTGTGAGACCAAACGTTGACACTAGTG |
| | Glyma07g14530-F2 | CCCTTGTTTCTATCCCAGCTTTC |
| | Glyma07g14530-R2 | GTCCTCGAAGTACACTGAGGG |
| | Glyma07g14530-R3 | CTTCCAATTCCATGAAGCTATTG |
| **Glyma07g14470 primers for cloning in arabidopsis** | | |
| | Gm470-Phy-F<br>Gm470-R | GGGGACAAGTTTGTACAAAAAAGCAGGCTTCGTTGTGACCGTCGTGGGTT<br>GGGGACCACTTTGTACAAGAAAGCTGGGTGAACAAATTGCAGGGTTGCAGAAATG |
| **qRT-PCR primers** | | |
| Glyma07g14530 (PCR efficiency = 1.93) | 530qRT-F6<br>530qRT-R6 | CGTTGCCAAAGATACCGTAGTGC<br>GGATACACAGGAGGGTAGCTAC |
| Metalloprotease (PCR efficiency = 1.76) | Metalloprotease-F<br>Metalloprotease-R | ATGAATGACGGTTCCCATGTA<br>GGCATTAAGGCAGCTCACTCT |
| Transgene terminator (PCR efficiency = 1.71) | qRbcSt-F<br>qRbcSt-R | GTTCGAGTATTATGGCATTGGGAAAACTG<br>CACAGTTCGATAGCGAAAACCGAAT |
| PR10 (PCR efficiency = 1.86) | PR10-F<br>PR10-R | AGTTACAGATGCCGACAACG<br>CCTCAATGGCCTTGAAGAGA |
| **Cloning primers** | | |
| GmUbiP:Glyma530-R:rbcST and GmUbiP:Glyma530-F:rbcST | *Asc*I-530-OE-F<br>*Avr*II 530-OE-R | ATTA**GGCGCGCC**ATGGAATCAGCGGCAAGAACA<br>CACG**CCTAGG**GTCAGCAAGTAGGACGCAAAG |

| | | |
|---|---|---|
| GmUbiP:1510-530:rbcST | *Asc*I-1510-530targ-F | ATATA**GGCGCGCC***AGGTGGAATAGGAAAAACAACT*GTGTACTTCGAGGACCTAAAC |
| | *Avr*II-530targ-R | TATAT**CCTAGG**ATCTTAGGGTTTCCCTAACGG |
| **Primers for confirmation of transgenic events** | | |
| StUbi3P:*hpt*:StUbi3T | Hygro117-F | CGATGTAGGAGGGCGTGGATA |
| | Hygro504-R | GTCGTCCATCACAGTTTGCCA |
| GmUbiP:Glyma530-R:rbcST and GmUbiP:Glyma530-F:rbcST and | Gmubi842-F | CGAGATTGCTTCAGATCCGTA |
| GmUbiP:1510-530:rbcST | RbcSt110-R | CCATTTCCATTTCACAGTTCG |
